# HIV IMPAIRS AND EXPLOITS PULMONARY TH17 AND TH22 CELL-MEDIATED IMMUNE RESPONSES TO *MYCOBACTERIUM TUBERCULOSIS*

**DOI:** 10.1101/2025.04.16.649152

**Authors:** Yazmin B. Martinez-Martinez, Matthew B. Huante, Kubra F. Naqvi, Mithil N. Shah, Joshua G. Lisinicchia, Megan A. Files, Jaid Perez, Benjamin B. Gelman, Mark A. Endsley, Janice J. Endsley

## Abstract

Tuberculosis (TB) kills an estimated 1.25 million people annually and is the leading cause of death in people with HIV (PWH) (1). The CD4^+^ T helper (Th) populations play significant roles in protective immunity to *Mycobacterium tuberculosis* (Mtb) and are essential hosts for HIV pathogenesis. Emerging evidence in blood and gastrointestinal mucosa of PWH suggests that, among Th cells, Th17 and Th22 may be preferentially depleted during HIV infection. Targeting of Th17 and Th22 cells by HIV could pose important and poorly understood risks for Mtb containment in those with co-infection. Mtb-driven activation of Th17 and Th22 immunity may also contribute to HIV proliferation and persistence. We employed a humanized mouse model of co-infection to assess changes in Th17 and Th22 frequency and function due to infection with HIV, Mtb, or both. In infected mice, Th17 cells were the predominant host for HIV in spleen and shown to be a source of HIV replication in pulmonary TB granulomas. Th17 cells were increased in lung of mice with TB or TB-HIV. Conversely, Th22 cells were reduced in mice with HIV or TB-HIV. Mtb infection increased the viral load in lung of co-infected mice while HIV suppressed the pulmonary Th17 family cytokine response to Mtb including IL-6, IL-22, IL-23, and IL-1β. Differential transcriptome assessment demonstrated that HIV co-infection disrupted Th17 pathways activated by Mtb in lung. Overall, these results suggest that HIV may compromise Th22 immunity and exploit Th17 cells to promote viral pathogenesis in the setting of Mtb and HIV co-infection.

## 1. Introduction

Tuberculosis (TB), caused by the *Mycobacterium tuberculosis* complex (Mtb), kills 1.25 million people throughout the world every year, including 161,000 people infected with HIV (PWH)(1). One-third of HIV deaths are due to TB, and the risk of developing active TB increases markedly in PWH, especially if latent tuberculosis is present (2). Addressing health problems resulting from Mtb and HIV co-infection is challenging due to the limitations of experimental models and the poorly understood microbial synergy that exacerbates both diseases (3). CD4+ T cells (T helper cells, Th) are critical in protection against TB (4, 5) but are also the main replication targets for HIV. Th cells in the lung, especially lung interstitium, may be lost early during Mtb-HIV co-infection, even before systemic immune impairment, and residual impairments in Th cells persist despite anti-retroviral therapy (6).

The effects of HIV or SIV infection on the numbers and function of Th1 and Th2 subpopulations have been characterized in accessible human specimens and tissues of NHP (non-human primate) models, including Mtb co-infections (7, 8). In contrast, less is known about effects on other Th subpopulations such as Th17 and Th22 cells, especially in organs of co-infection, such as lung and spleen. Th17 cells promote inflammation and mediate protective immune responses to various bacterial and fungal pathogens (9–11). These beneficial responses of Th17 occur mainly at mucosal barriers (12, 13) through secretion of cytokines such as IL-17A, IL-17F, IL-22, and IL-21, and by neutrophil recruitment (14, 15) in gut and lung mucosa (13, 16). Th22 cells secrete IL-22 but not IL-17. IL-22 targets epithelial cells, fibroblasts, and macrophages specifically involved in tissue repair, wound healing, and mucosal immunity (17). Th17 are important in vaccine-induced protection against Mtb and strongly correlate with bacterial control in mouse and humans, while Th22 cells and their cytokines contribute to immunity through incompletely described mechanisms (18–25). During active TB, both Th17 and Th22 are increased in lung but decreased in the blood (26–29).

Understanding the multifaceted effects of HIV on diverse T-cell populations is a challenge for vaccination and other host directed interventions. The basis for Simian Immunodeficiency Virus (SIV)-mediated reactivation of latent TB is associated with CD4+T cell loss, changes in T cell cytokine responses, and CD4+T cell loss-independent mechanisms in different NHP species (7, 30, 31). During HIV mono-infection, lack of Th17 is linked to impairment in the Th17/Treg ratio (32) and disease progression (33), where memory Th17 (CD45RA-CCR6+) are preferentially infected and depleted (34–36). This preferential depletion of Th17 among Th cells is due to higher levels of CD4, CCR5, and CXCR4 (receptors and co-receptors for HIV) and a lack of anti-HIV-RNases (37, 38). IL-22 is protective against HIV-induced gut epithelial damage (39). However, both Th17 and Th22 are depleted in blood, sigmoid colon, jejunum, and colorectal mucosa following infection with HIV or SIV (39–42). Mtb specific Th17 and Th22 cells, as well as IL-17 and IL-22 cytokines, are decreased in blood of Mtb-HIV co-infected subjects (43–46).

Collectively, observations from PWH and NHP models of SIV suggest that HIV may impair protective roles of Th17 and Th22 against Mtb in co-infections. However, experimental validation and extension to critical tissue sites of infection are limited to date. The human immune system (HIS) mouse model has gained acceptance due to the development of functional human immune compartments capable of reproducing important human immune responses to pathogens with restricted host range. Benefits of HIS mice compared to NHP models are the smaller animal size, the ability to use HIV unmutated strains, reduced regulatory burden, maintenance cost, and scalable sample size for statistical analysis (47). HIS mouse models have demonstrated reliable aspects of human infection and disease progression in HIV and Mtb-HIV co-infection (48–51) as well as translational utility for assessment of host directed therapies in these diseases (52, 53).

Here, using a cord-blood HIS infection model, we demonstrate that Th17 cells in tissues are the predominant host Th cell for HIV. Viral load was increased in mice with pulmonary TB compared to mice with HIV only, and RNAScope analysis identified cells co-expressing IL-17 and HIV *gag* transcripts in TB granulomas. Th17 cells remain elevated in lung of mice with Mtb, or Mtb and HIV co-infection, while activation of Th1 cells was suppressed in co-infection. Ingenuity Pathway Analysis of bulk RNASeq data from lung of Mtb-infected mice predicts disruption of IL-17 signaling pathways due to HIV co-infection. We further demonstrate that HIV markedly suppressed Th22 responses in both HIV mono- or co-infected mice. Overall, these findings indicate that HIV could exploit Th17 cells to promote viral pathogenesis in the setting of Mtb-HIV co-infection, leading to an increased viral reservoir at important sites of immunity in the lung and further suppression of immunity mediated by events downstream of IL-17 receptor signaling pathways. Th22 protection against TB could be lost early during co-infection due to the generalized suppression of Th22 cells and IL-22 cytokine following HIV infection. Preventive and therapeutic interventions for TB should thus consider these effects of HIV on Th17 and Th22 responses (**Fig 1**) in Mtb-HIV co-infected individuals.

**Fig 1.**
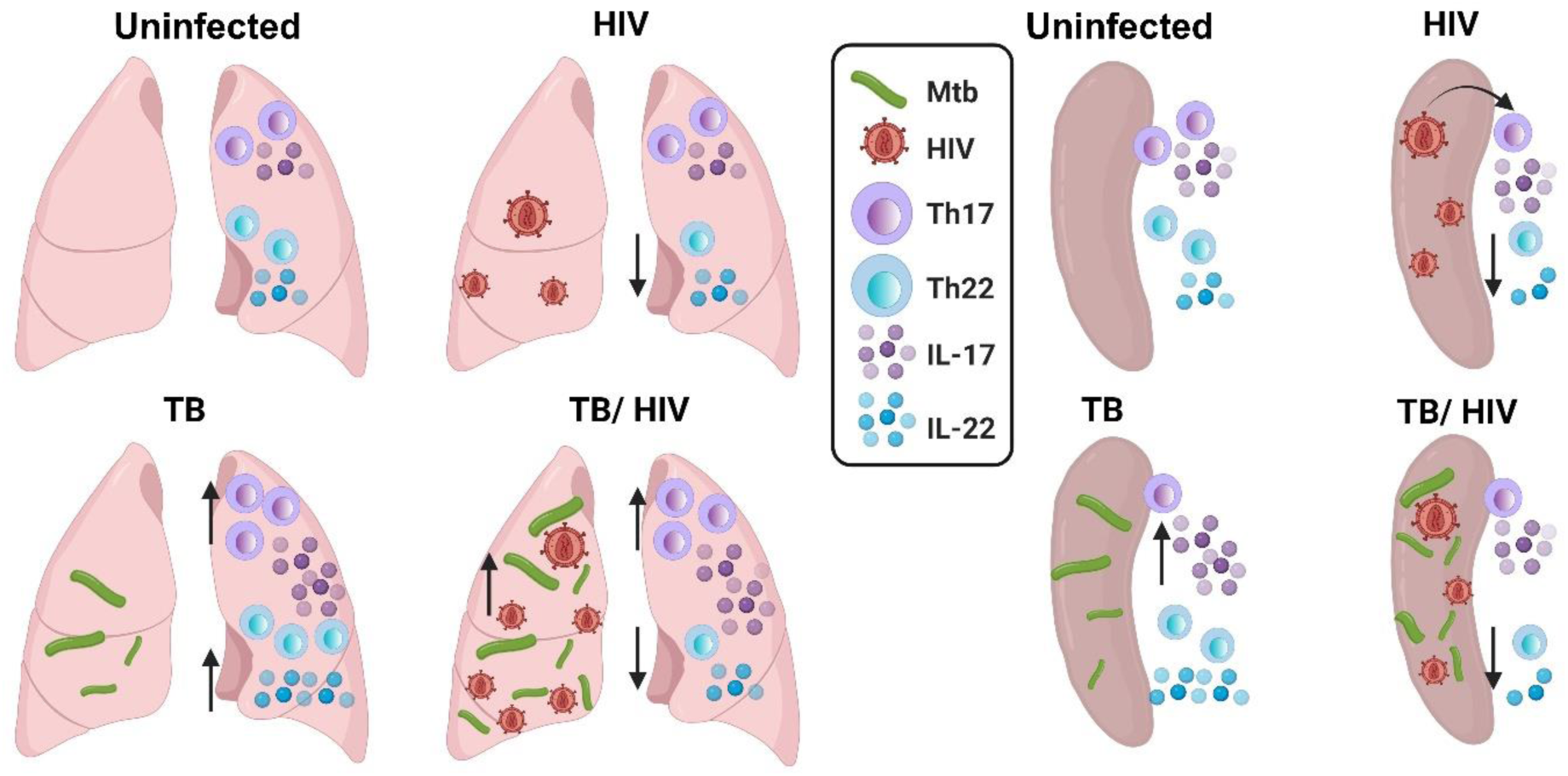
Graphical abstract of Th17 and Th22 impairments by HIV and Mtb-HIV co-infection in lung and spleen. Graphical representation of Th17 and Th22 responses in the lung (left section) and spleen (right section) in the four conditions of each panel: uninfected (upper left), HIV (upper right), TB (lower left) and TB-HIV (lower right). In the lung there is increased HIV viral load in TB-HIV compared to HIV, increased Th17 and IL-17A when there is infection with Mtb (TB and TB-HIV), and decreased Th22/IL-22 when there is HIV infection (HIV and TB-HIV). In spleen, Th17 are preferentially infected by HIV during HIV mono-infection, IL-17 cytokine is increased during TB, and Th22/IL-22 are impaired when there is HIV infection (HIV and TB-HIV groups). Image created with BioRender.com.

## 2. Materials and Methods

### HIS mouse infection models

All animal experiments were approved under protocol 1501001B, complying with the University of Texas Medical Branch (UTMB) Institutional Animal Care and Use Committee. The NSG-hu-CD34+ HIS mouse model was used for both animal experiments **(Fig 2A** and **3A)**. NOD-SCID/γc^null^ female mice (strain 005557) were purchased from The Jackson Laboratory (JAX) following injection of with human CD34^+^ stem cells via an i.v. (intravenous) route and used at 36 weeks of age. For both experimental timepoints, human T-cell reconstitution was validated in peripheral blood via flow cytometry using anti-CD45 human-AmCyan, anti-CD45 mouse-PE, and anti-CD3 human-BUV395.

**Fig 2.**
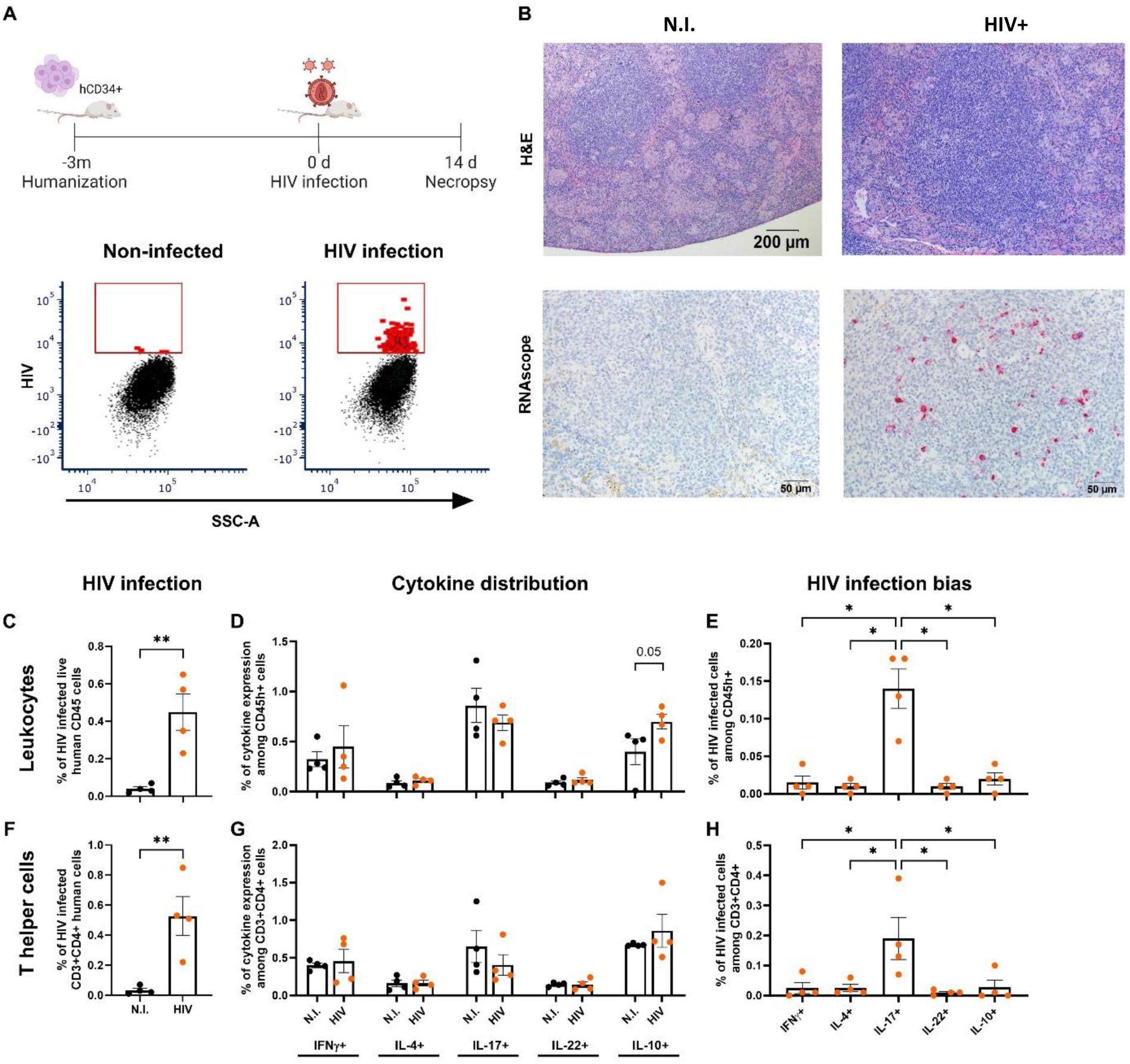
Th17 and IL-17+ leukocytes in spleen are preferentially hosts for HIV. (A) Experimental design to generate HIS mice and assess HIV distribution among Th subpopulations and representative flow cytometry analysis. Immune-deficient NOD.Cg-*Prkdc^scid^ Il2rg^tm1Wjl^*/SzJ (NSG) mice were reconstituted with human immune cells by i.v. administration of cord blood CD34+ stem cells. After confirmation of human immune system reconstitution, mice were infected i.v. with 10^5 HIV-1 ADA TCID, and spleens collected after 14 days of infection for flow cytometry and histology analysis. Representative examples of flow cytometry gating to assess HIV-FITC-A signal in non-infected (N.I., left) and HIV-infected (right) HIS mice. (B) Representative images of hematoxylin and eosin (H&E, top) and RNAscope-based detection of HIV gag (bottom, red substrate) in comparison between non-infected (left) and HIV-infected (right) spleen. (C-H) Preferential infection of Th17 and IL-17+ leukocytes in comparison to other cytokine producing cell subsets present in the spleen at 14 days post HIV infection. (C, F) Increase in intracellular signal for HIVp24 in leukocytes and CD3+CD4+T cells of HIV-infected animal spleen compared to background fluorescent signal in non-infected animals. (C-E) Leukocytes and (F-H) T helper (CD3+CD4+) cell infection with HIV. (D, G) Similar proportions of cytokines (IL-4, IL-10, IL-17, IL-22 and IFNγ) are produced by total leukocytes and Th cells from spleen of uninfected and infected animals. (E, H) HIV infection bias towards IL-17+ leukocytes and Th17 in comparison to other cytokine-producing cells at 14 dpi. n=4/group. Differences between two groups were determined using a Student’s T-test while an ANOVA with the Benjamini FDR post-hoc analysis was used for analysis of data from three or more groups. Significance was considered for *p<0.05 **p<0.01.

**Fig 3.**
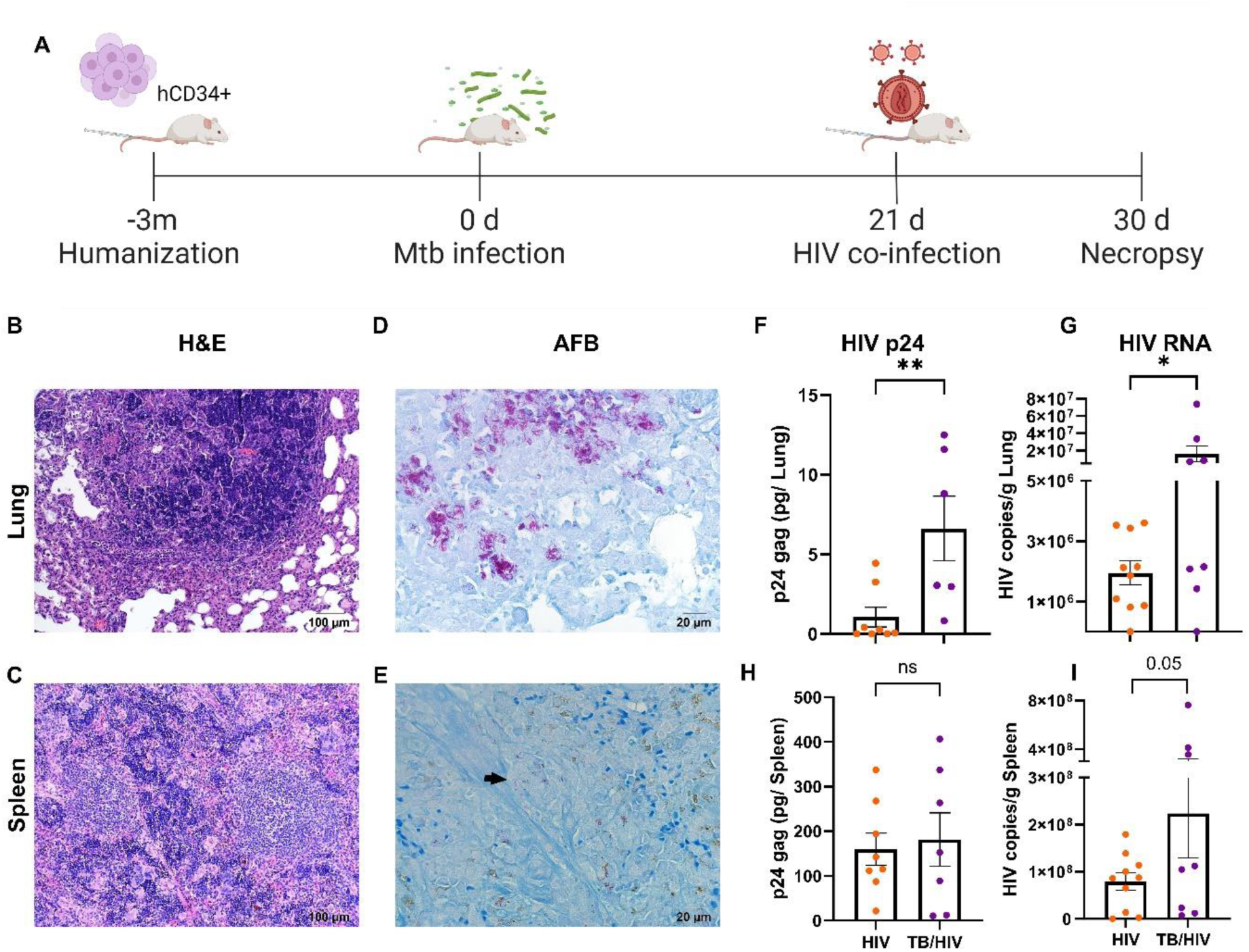
Mtb HN878 infection increases HIV replication in lungs of co-infected HIS mice. (A) Experimental design to generate acute Mtb-HIV co-infection in HIS. After confirmation of human immune system reconstitution, mice were infected with 100 CFU Mtb HN878 via an aerosol route for a total of 30 days. At day 21 p.i., non- and Mtb-infected groups were further infected i.v. with HIV-1 ADA (10^5^ TCID) to establish 4 groups: non-infected, HIV, TB, and HIV-TB. The study included N=38 mice with n=8-10 / group. Image created with BioRender.com. (B, C) Representative images of lung (B) and spleen (C) tissue specimens showing granulomatous inflammation from Mtb-HIV co-infected tissues as visualized with hematoxylin and eosin (H&E) (D-E). Representative lung (D), or spleen (E) from Mtb-HIV co-infected HIS mice infected with Mtb bacilli, as visualized with AFB staining. Arrows indicate Mtb bacilli clusters. (F-I). HIV viral load as determined by ELISA-based detection of HIV p24 in lung (F), and spleen (H). HIV viral copies per gram of lung (G) and spleen (I) in mice infected with HIV or HIV-Mtb were determined by using RT-PCR. Differences between the two groups was determined by a Student’s T-test with significance considered for *p<0.05, **p<0.01, and ns= non-significant.

All animal and bacterial experiments were performed in Biosafety level 3 (BSL3) laboratory and animal facilities (ABSL3) approved by the Centers for Disease Control and Prevention. Experiments were conducted under established guidelines and protocols from the UTMB Department of Biosafety. Mtb strain HN878 (Beijing lineage) was cultured to mid-log phase for delivery of aerosol infection with 100 CFU/mouse as previously described (29). HIV-ADA (NIH AIDS Reagent Program) was propagated in human PBMCs and concentrated by Lenti-virus concentrator (Origene, Cat. TR30025) for animal injections. Virus stock was quantified by GHOST cell infections, measuring the Tissue culture infectious dose (TCID) by GFP quantification via flow cytometry, as described previously (54). Virus stock was diluted with PBS to 1×10^6^ TCID/mL, and further HIV infections were performed via tail vein injection with 100 µL containing 100,000 TCID/ mouse.

For the HIV-mono-infection experiment **(Fig 2A)**, mice were infected with HIV for 14 days, or non-infected, and humanely euthanized to collect spleen for histology and flow cytometry (n=4 per group).

For the acute Mtb-HIV co-infection **(Fig 3A)** mice were infected by the aerosol route with Mtb, or non-infected, as previously described (29). At day 21, non- and Mtb-infected groups were further i.v. infected with HIV-1 (strain ADA) as described (53). All mice were humanely euthanized at experimental day 30, establishing four experimental groups: non-infected, HIV (9 days), Mtb (30 days), and Mtb-HIV (30 days Mtb infection with HIV co-infection during the last 9 days) with 8-10 mice/group (N=38). Lung, spleen, blood, and liver were collected for further assessment.

### Assessment of cell populations by flow cytometry

The left lung lobe and one-third of the spleen were collected for flow cytometric analysis. Tissue was disrupted, supernatants were collected for cytokine measurements, and the cells were activated with human anti-CD3, anti-CD28, and GolgiStop (BD) to retain intracellular cytokines, as previously described (29). Extracellular staining was performed with live/dead Near IR and antibodies specific to human markers: CD3-BUV395, CD4-PacBlue, CD45-AmCyan, and CD14-BV786. After fixation with Cytofix/Cytoperm, intracellular staining was performed with antibodies to human cytokines IFNγ-BV650, IL-10-AF594, IL-17A-PE, IL-22-APC, CD68-PerCP-Cy5.5, and to HIV-FITC. Fixation was performed with 4% ultrapure formaldehyde (FA) for 48 hours to ensure inactivation, followed by a change to 1% FA and acquisition of total cells on an LSR II (Fortessa) flow cytometer. Subsequent analysis was performed using FCS Express 6 software (de Novo, Inc).

### Determination of viral burden

Quantification of HIV p24 capsid protein was used to estimate viral burden in lung, and spleen extracellular supernatants, using an ELISA kit (Zeptometrix, Cat No. 0801111).

To evaluate HIV RNA levels, the right inferior and post-caval lobes from lung, one-third of spleen, or a section of the left superior lobe from liver were collected. RNA extraction was performed using the Qiagen RNeasy mini kit (Cat. 74104), and the RNase-free DNase set (Cat. 79254), according to the manufacturer’s instructions. This RNA was used for assessment of viral copies by measuring the HIV gag mRNA per gram of tissue, detected by RT-PCR as previously described (55).

### Measurement of human cytokines

Extracellular cytokine measurements were done following the collection of intracardiac blood and centrifugation at 3,000 rpm for 10 minutes. Supernatants from lung and spleen were also assessed following tissue disaggregation and centrifugation at 350 ×g for 5 min before cellular activation. Lung, blood, and spleen samples were collected in cryovials and frozen at −80°C for subsequent analysis. Frozen samples were inactivated by γ-irradiation on dry ice for biosafety as described (56) following approved UTMB biosafety protocols. The human cytokines from lung and spleen were diluted as they were extracellular, without cell activation, and obtained from tissues of a xenochimeric mouse model. For these reasons, we concentrated the protein levels in lung and spleen supernatants and incubated them overnight in the detection system for higher detection. The six most representative animals from each group were selected, and the supernatants were concentrated using the Pierce™ Protein Concentrator PES, 3K MWCO (Cat 88512). Human cytokines from plasma or concentrated lung and spleen samples were quantified using a Bio-rad multiplex ELISA, the Bio-Plex Pro Human Th17 cytokine assays (Catalog #: 171AA001M), following the manufacturer’s instructions, except for incubation of samples with beads overnight. IL-21 and IL-31 results from the kit were excluded, as they were below the detection limit.

### Histopathology and RNAscope

The right superior and middle lobes from lung were inflated with and collected in 10% Neutral Buffered Formaldehyde (NBF) containers (No. 032-059) along with one-third of the spleen and the left superior lobe of the liver. After 48h and a formalin change, tissues were removed from BSL3 and sent to the Anatomic Pathology Laboratory at UTMB. Paraffin-embedded tissue sections were serially sectioned (5 µm) and stained at the same core facility with H&E and the Ziehl-Neelsen method to detect acid-fast bacilli (AFB).

Additional tissue sections were stained using *in situ* hybridization to detect HIV-viral infection in Th17 cells or other T helpers surrounding the TB-granuloma. RNAscope 2.5 HD chromogenic-RED assay for single-plex HIV RNA detection, or RNAscope Multiplex Fluorescent reagent kit v2 assay (Advanced Cell Diagnostics, ACD Cat. 323100) were used following the manufacturer’s instructions. Targets detected were HIV RNA-gag-pol (probe 317691) for single-plex analysis. HIV RNA (non-gag-pol probe 317711-C2), human IL-17 RNA (probe 310931-C3), and human CD4 RNA (probe 605601) were detected for multiplex analysis. Imaging was later performed by the UTMB Optical Microscopy Core, with the confocal microscope Zeiss LSM 880 with Zen software, followed by the program Fiji.

### Differential transcriptome analysis

High throughput RNA sequencing was performed for 24 HIS mouse lung samples from six replicates of four conditions (non-infected, Mtb, HIV, or Mtb-HIV). The samples selected for RNASeq were from the same tissues used to assess cytokine levels and viral burden. RNA quality was assessed with an Agilent Bioanalyzer, and RIN (RNA Integrity Number) values ranged between 9.8 and 10. PolyA+ RNA was purified from ∼100 ng of total RNA. NEBNext Ultra II RNA library kit (New England Biolabs) was used to prepare the sequencing libraries following the manufacturer’s protocol. Libraries (twenty-four) were quantified by qPCR, pooled, and sequenced using a single-end 75 base run on a Next-Seq 550 High-output flow cell at the UTMB Next Generation Sequencing Core. Canonical Pathway studies and IL-17 pathway disruption analyses were performed by Ingenuity Pathway Analysis Software (IPA, Qiagen).

### Statistical analysis

Statistical analysis and graphical presentations were developed using GraphPad Prism 10. software. Data is presented as the mean ± SEM. For comparison between two experimental groups, an unpaired T-test was used. One-way ANOVA was used for experiments with more than two groups, with post-hoc correction for multiple comparisons. The false discovery rate was controlled by using the two-stage test set-up method for Benjamini, Krieger, and Yekutieli. Significance was considered with a p-value <0.05, and a trend in significance was considered where p<0.1.

## 3. Results

### HIV preferentially infects Th17 cells in HIS mouse tissue

In PWH, peripheral Th17 appear to be a preferred Th cell target for HIV infection and may act as a viral reservoir in blood and colorectal mucosa (40, 57, 58). To validate and extend these observations in the HIS mouse, we assessed cellular distribution of HIV in Th populations using the spleen as a source of peripheral leukocytes. HIV infection was confirmed in the spleen at 14 days post-infection based on flow cytometric detection of intracellular HIV p24 protein **(Fig 2A, Sup. Fig 1B)** and RNAscope-based detection of viral RNA **(Fig 2B)**. HIV infection was detected in human CD45+ cells (leukocytes) as depicted in **Figure 2C**. The percentage of leukocytes producing specific cytokines (IFN-γ, IL-4, IL-17, IL-22, and IL-10) did not significantly differ among non-infected and HIV-infected tissues (**Fig 2D**), indicating a lack of subset-specific depletion at this acute infection stage. A definitive bias for HIV infection of the IL-17+ leukocyte subset, however, was observed (**Fig 2E**). Analysis of the Th subpopulations (CD3+CD4+) demonstrated a highly similar pattern of HIV infection, cytokine production, and HIV infection bias among the cytokine-producing cells that generally define Th subpopulations (**Fig 2F, G, and H**). These results demonstrate preferential infection of IL-17-producing leukocytes and Th17 cells by HIV in the spleen.

### Co-infection with Mtb promotes an increase in pulmonary HIV replication

To demonstrate the clinical relevance of the model for HIV infection in the setting of established TB, HIS mice were infected with Mtb and subsequently infected with HIV. Before infections, reconstituted mice were distributed in the four experimental groups normalized for human leukocyte reconstitution **(Sup. Fig 2)**. Reconstitution average percentages in mice were 57% human among total leukocytes **(Sup. Fig 2A)** and 67% of human T cells among the human leukocytes **(Sup. Fig 2B)**. Mice were infected with 100 CFU Mtb HN878/mouse via an aerosol route, for 30 days. On day 21, non- and Mtb-infected groups were further infected with HIV-1 ADA via an i.v. route to establish four groups: non-infected, HIV, TB, and TB-HIV, as described in **Figure 3A**. Assessment of tissue histology demonstrated development of organized granulomatous inflammation in lung, spleen (**Fig 3B, C**), and liver (**Sup. Fig 3A**). Abundant acid-fast bacilli (AFB) were detected throughout lesions and central areas of inflammation in lung, liver, and spleen (**Fig 3D, E, Sup. Fig 3B**). Analysis of HIV infection of tissues by using ELISA and RT-PCR revealed greater HIVp24 protein in lung supernatants (**Fig 3F**) and increased viral RNA copies in lung of mice with Mtb co-infection (**Fig 3G**). The splenic viral burden was markedly increased compared to the lung in general (**Fig 3F-I**). However, co-infection with Mtb did not result in significant changes to viral burden in the spleen and liver, although a trend (p=0.05) towards increased viral transcription in spleen was observed (**Fig 3I, Sup. Fig 3C**).

### HIV co-infection promotes distinct shifts in pulmonary T cells and macrophage responses to Mtb infection

To determine the effect of acute HIV, Mtb, or HIV/Mtb infections on important immune populations in the lung and spleen, we performed flow cytometric analyses of single-cell suspensions from disrupted tissue. The representative flow cytometry gating is shown in **Sup. Fig 1A**. In the lung, there were fewer total human leukocytes in HIV and TB groups, compared to non-infected, and interestingly, to TB-HIV **(Fig 4A)**. Among lung leukocytes, there were fewer T cells as a percentage in the TB and TB-HIV, compared to HIV and non-infected groups. These effects in the TB and TB-HIV groups were independent of CD4+T cell loss (**Fig 4A**), despite the increased viral load measured in the lung (**Fig 3F, G**). In contrast, a moderate and non-significant decrease in Th cells in the HIV-infected group corresponded with decreased leukocytes (**Fig 4A**). The proportional decrease in T cells was compensated by increased macrophages in TB group. Interestingly, the percentage of macrophages in lung of the TB-HIV group remained similar to non-infected and HIV groups. This indicates that changes in another cell population may contribute to the shifts in % of T cells in this group **(Fig 4A)**. In contrast to the lung, the spleen did not present changes in the leukocyte populations, as observed in **Fig 4B** although HIV groups were generally, but non-significantly, reduced.

**Fig 4.**
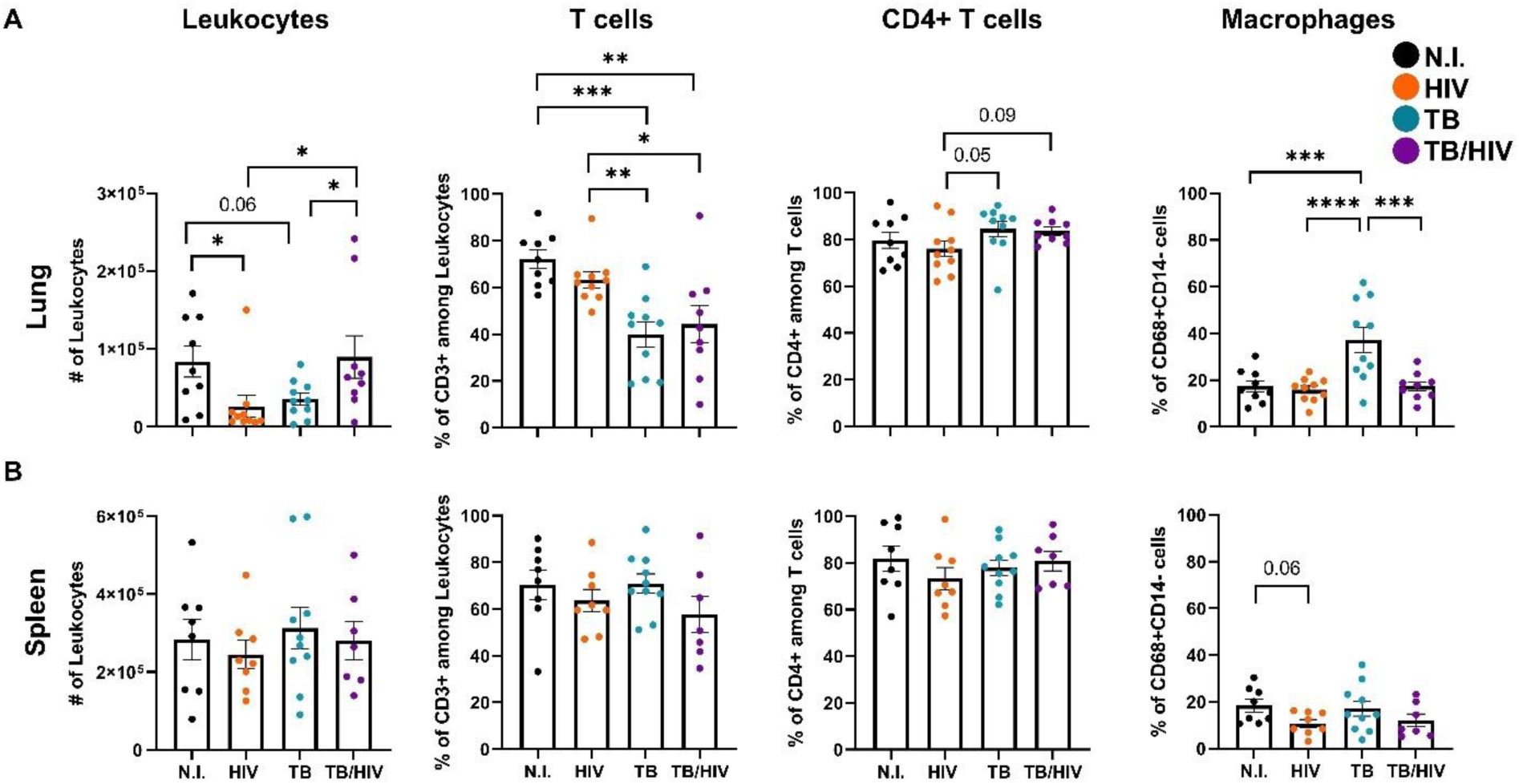
Co-infection promotes a differential lung Macrophage and T cell profile compared to Mtb or HIV mono-infection. Comparison of lung (A) and spleen (B) immune populations in non-infected (N.I.), HIV, TB, and TB-HIV groups. Total leukocytes among viable singlet cells were selected based on detection of human CD45 as shown in Supplemental Figure 1. Among human CD45+ cells, T cells and macrophages were identified by surface expression of CD3 (Total T cells), CD3 and CD4 (Th cells) and CD68+CD14lo (macrophages) and shown as % of human CD45 cells. Differences were determined by using one way ANOVA with the Benjamini FDR post-hoc used to determine differences among different groups. Significance was considered for *p<0.05, **p<0.01., ***p<0.001, ****p<0.0001

### The Th17 response to pulmonary TB differs from Th1 and Th22 in the setting of HIV co-infection

Th1, Th17, and Th22 are expanded in the lung during active TB in human and murine models (27–29). Different responses have been described in the spleen: Th17 cells were shown to decrease while Th IL-10+ increased during active TB in a murine model (29). To determine how HIV co-infection affects the Th populations (Th1, Th17, Th22, and ThIL10+) in lung during Mtb-HIV co-infection, we assessed changes in functional Th subsets and the relative quantity of cytokine produced by flow cytometry and Bioplex analysis **(Fig 5 A-F)**. As expected, Th1 percentages and extracellular IFNγ were increased in the TB group **(Fig 5A)**, consistent with changes observed in human lung. However, expansion of Th1 due to Mtb infection is suppressed by HIV infection in accordance with Geldmacher’s findings in human subjects (59). A similar decrease was observed for soluble IL-10 without changes in ThIL-10+ cells **(Sup. Fig 4F)**. Interestingly, human Th17 and IL-17 cytokine increased in lungs of TB and remained elevated in the co-infected group **(Fig 5C)**. In contrast, Th22 cells and IL-22 cytokine production were suppressed by HIV in the mono- and co-infected groups (**Fig 5E).**

**Fig 5.**
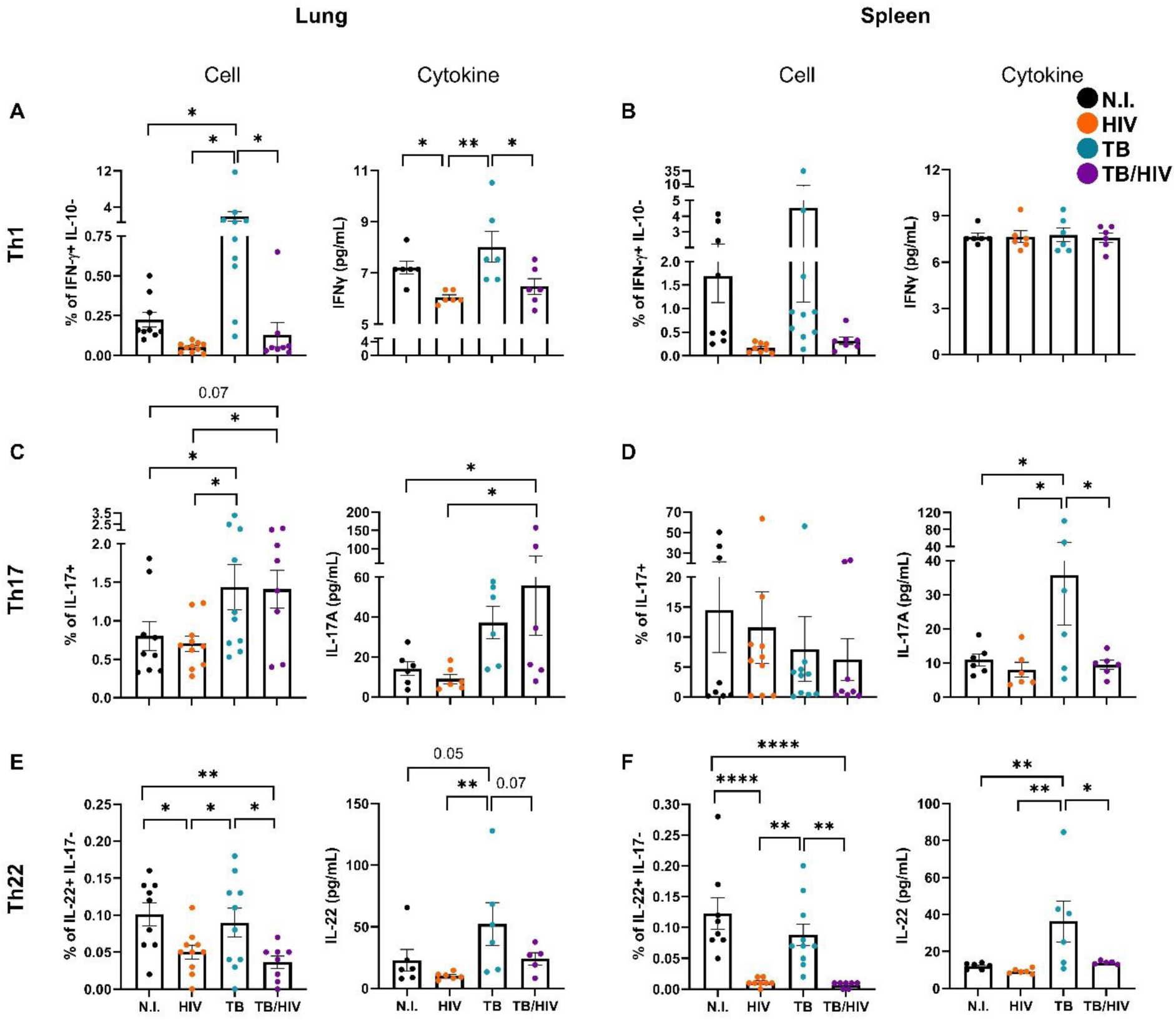
Th17 and Th22 cell defects in co-infected mice reveal tissue and subpopulation effects. T helper subset comparison in lung (A, C, E) and spleen (B, D, F) between the four experimental groups: non-infected (N.I.), HIV, TB, and TB-HIV. For each organ (lung or spleen) subsection, the left panel represents the cell population, and the right panel shows the respective cytokine in tissue supernatant. Flow cytometry was used to identify Th subpopulations based on intracellular cytokine staining for IL-17, IL-22, and IFNγ, or IL-10 in viable CD45+CD3+CD4+ T cells using gating strategy shown in Supplemental Figure 1. Th22 cells were defined as those with an IL-22+ and IL-17-phenotype. Soluble cytokines from lung and spleen were measured by Bioplex and are depicted in pg/mL of organ supernatant. Significance was considered for *p<0.05, **p<0.01, ***p<0.001, ****p<0.0001 following One-Way ANOVA and a Benjamini FDR correction for multiple comparisons.

Analysis of the spleen as a peripheral lymphoid compartment revealed a more variable and subdued response overall compared to the lung. Th17 cells were markedly more abundant in the spleen as a percentage of total T cells irrespective of infection status. No significant difference in Th1 cell or soluble IFNγ production was observed **(Fig. 5B).** A unique observation in the spleen was the HIV-mediated suppression of IL-17 cytokine production in response to Mtb that occurred in the absence of changes in Th17 cells (**Fig 5D**). The suppressive effect of HIV on Th22 and IL-22 was more pronounced in the spleen compared to the lung, HIV mono-infection was also associated with increased ThIL-10+ cells in the spleen (**Sup**. **Fig 4G**). These outcomes may reflect the increased viral burden in spleen compared to lung as we observed in **Fig 3**.

### Cytokine determinants of Th17 and Th22 subset bias are differentially activated in Mtb and Mtb-HIV infections

To determine immune drivers of the unique subset activation and suppression in Mtb and HIV infections, we assessed a panel of cytokines and other soluble mediators (Bio-Plex Pro Human Th17 cytokine assay) that are key to Th17 and Th22 differentiation (17, 60) across the lung, spleen, and blood compartments **(Fig 6, Sup. Fig 4 and 5)**. IL-1β and IL-6 were increased in response to Mtb infection across all compartments while a consistent suppressive effect of HIV co-infection was observed **(Fig 6A)**. Of note, some effects were less robust (p=0.05-0.07) due to animal variation. Activation of IL-23 due to Mtb infection was markedly suppressed due to HIV co-infection in lung but was otherwise similar among treatment groups in the spleen and plasma (**Fig 6C**). The IL-17F response was similar to IL-1β and IL-6 across compartments except that plasma responses were low and did not differ among groups (**Fig 6D**). Mtb infection also increased IL-4 and the alarmins IL-33 and IL-25 (IL-17E) in the lung and IL-4 in the spleen in agreement with previous reports of increased Th2 during TB (29) **(Sup. Fig 4)**. However, HIV co-infection demonstrated moderate suppressive effects of the Mtb activation response in these cytokines **(Sup. Fig 4)**.

**Fig 6.**
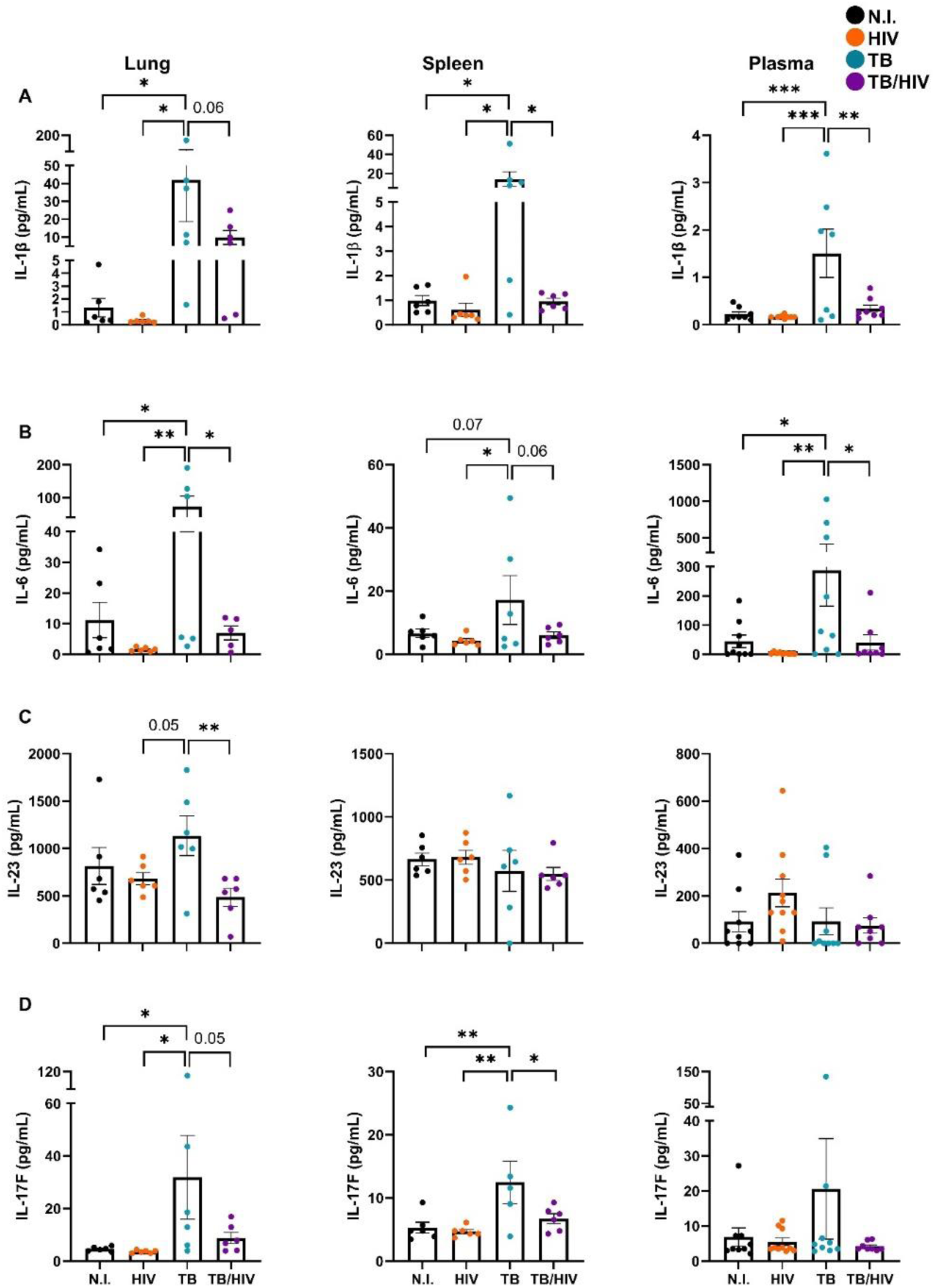
Th17 and Th22 cytokine pathways activated by Mtb are suppressed in Mtb-HIV co-infection. Comparison of extracellular human cytokines (pg/mL) from lung (left, n=6/group), spleen (center, n=6/group), or plasma (right, n=8-10/group) between the four experimental groups: N.I., HIV, TB, and TB-HIV. (A) IL-1β, (B) IL-6, (C) IL-23, and (D) IL-17F are increased due to Mtb infection, but decreased responses are observed after HIV co-infection. *p<0.05, **p<0.01, ***p<0.001, (One-Way ANOVA with Benjamini FDR correction).

These cumulative results demonstrate that HIV infection disrupts cytokine networks whereby Th subpopulation bias and functional activation is regulated. The effects are most pronounced at the site of Mtb infection (i.e. lung) but are measurable in the periphery, suggesting a more potent response occurring in the lung. Suppression of IL-1β by HIV **(Fig 6A)**, along with the moderate suppression of TNF that reached significance in plasma **(Sup. Fig 4, 5B)**, could be especially significant with regard to reduction of Th22 populations in TB-HIV **(Fig 5)** given the key roles of these two cytokines for Th22 phenotype differentiation (61–63).

### Th17, but not Th22, are important host cells for HIV in TB lung granulomas

Th17 are increased and protective during TB infection and are targets for tuberculosis vaccination efforts (64, 65). However, Mtb-specific Th17 responses are skewed by HIV in blood from HIV-chronic infected patients (66). To determine if Th17 are preferentially infected in lung or spleen during Mtb-HIV co-infection we analyzed the different HIV infection target cells. We observed that Th17 cells were uniquely resistant to HIV-mediated depletion in Mtb-infected HIS mouse lung, compared to Th1 and Th22 cells **(Fig 5C**). Since we had observed preferential infection of splenic Th17 by HIV **(Fig 2)**, we examined whether Th17 are an important cellular niche for HIV pathogenesis in TB granulomas. Detection of intracellular p24 viral capsid protein by flow cytometry revealed a similar distribution of HIV among Th1, Th17, and Th-IL-10+ cells in the lung of mice infected with HIV **(Fig 7A)**. Co-infection with Mtb did not significantly alter this pattern **(Fig 7B)**. The marked bias toward Th17 as the preferential host for HIV in the spleen that we initially observed **(Fig 2)** was reproduced in this set of mice reconstituted with separate donor stem cells **(Fig 7C)**. Following co-infection with Mtb, however, a highly variable and similar distribution of p24+ cells among Th1, Th17, and ThIL-10 subsets was observed **(Fig 7D)**. An especially interesting observation was the notable lack of detectable p24 in Th22 cells from the lung or spleen following HIV infection **(Fig 7 A, C)**. Co-infection with Mtb did not alter the susceptibility of Th22 cells to HIV infection in the lung or the spleen **(Fig 7 B, D)**.

**Fig 7.**
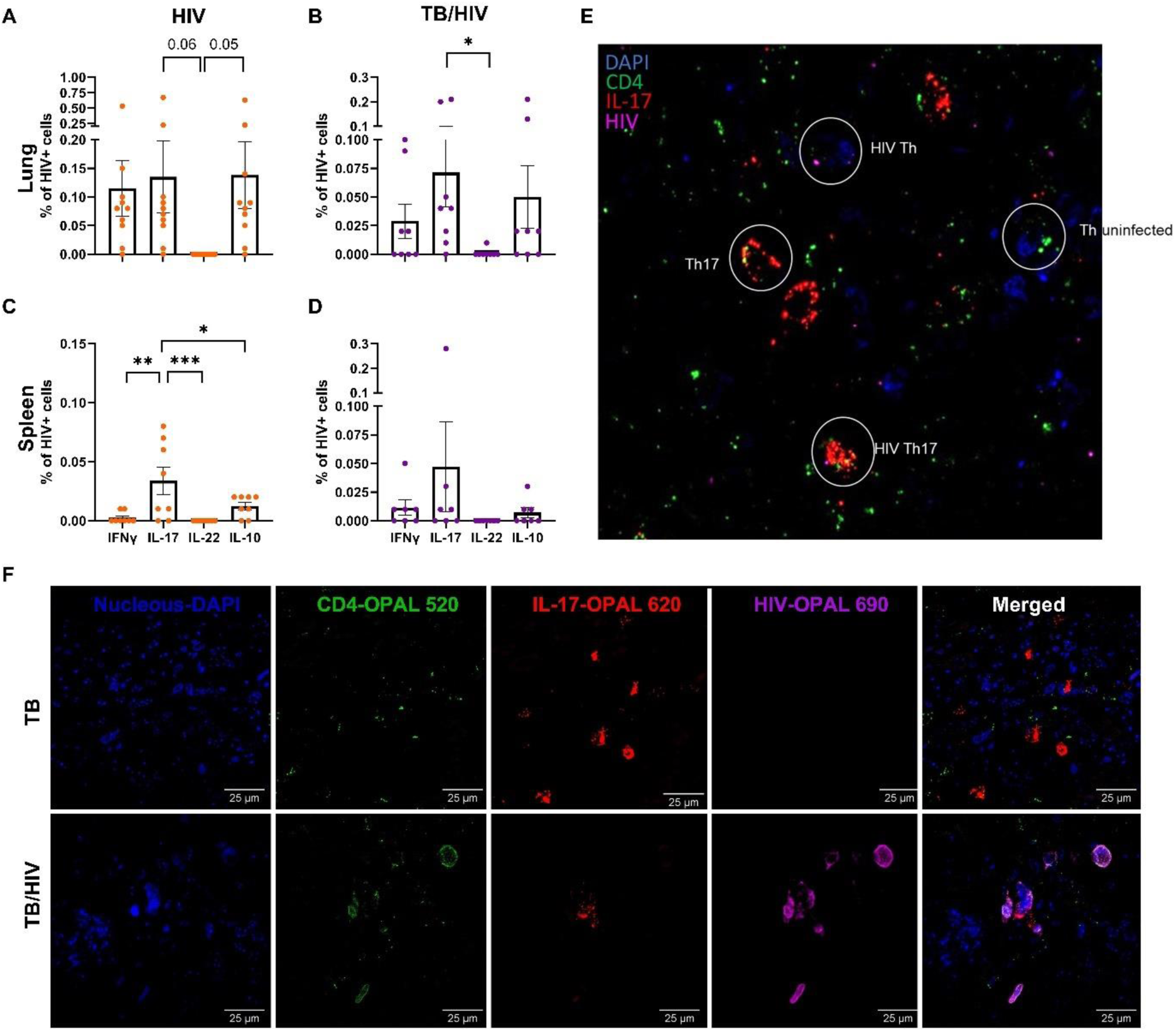
Th17 are preferential host in spleen and frequently found among HIV-infected cells in TB granulomas. (A-D) Percentage of HIV-infected cells expressing IL-17, IL-22, IFNγ, or IL-10 among T helper (viable CD45+CD3+CD4+ singlets) lymphocytes, in lung (A-B) or spleen (C-D), during acute (9 days) HIV infection (orange, A, C) or TB-HIV co-infection (purple, B, D). *p<0.05, **p<0.01, ***p<0.001 (One-Way ANOVA with Benjamini FDR correction). (E-F) Representative immunofluorescent RNAscope stained images, through *in situ* hybridization, detecting RNA of HIV (non-gag-pol) with OPAL 690 (purple), human IL-17 RNA paired with OPAL 620 (red), and human CD4 RNA with OPAL 520 (green). (E) Representative RNAscope image depicting possible combinations found in the lung TB-HIV granuloma. (F) Close-up of Th17 cells in lung TB lesions and infected by HIV in TB lesions of TB-HIV group, assessed by RNAscope.

HIV-infected cells surrounded granulomas (**Sup. Fig 6**) as previously shown (67), but also peribronchial areas, or blood vessels in the lung during Mtb-HIV co-infection. Th17 can be effectively infected, as shown in **Figures 7A-D**, and by RNAscope in **Fig 7E-F**. Even though human IL-17A is increased in both TB and TB-HIV groups, IL-17 transcripts were more evident in the TB group by RNAscope multiplex immunofluorescent imaging **(Fig 7F)**.

### Differential transcriptome analysis reveals HIV-mediated disruption of pathways that regulate the Th subpopulation response to Mtb infection in the lung

In order to identify the different pathways affected during Mtb-HIV co-infection that may cause upstream differentiation cytokine suppression (**Fig 6**), we performed high throughput bulk RNASeq of lung tissue (UTMB Molecular Genomics Core Facility) using six animals per experimental group. Differential transcription of human genes demonstrated both individual animal and experimental infection effects **(Sup. Fig 7A and Supplementary files)**. Ingenuity Pathway Analysis (IPA) of differentially expressed genes further identified the principal canonical pathways affected in the pertinent comparisons. Regarding the higher predicted scores for dysregulated pathways, IL-17 signaling pathways were present among the first affected groups **(Sup. Fig 7C)**. In support of the differences in Th populations and related cytokines that we observed (**Fig 5-6** and **Sup. Fig 4-5**) IPA identified the pathogen-induced cytokine storm signaling pathway as the immune network most impacted by HIV, especially the pathways activated by Mtb (**Sup. Fig 7C**). The IL-17 and IL-6 signaling networks were shown to be disrupted individually in a pattern similar to the overall cytokine storm pathway. Predicted disruptions of Th17 and Th2 pathways by HIV infection also reflects the potential for transcriptional regulation of cytokines and other molecules that determine Th subset differentiation and activation (**Sup. Fig 7C**). Another interesting finding is the oxidative phosphorylation pathway, predicted to be decreased in almost all comparisons, except TB-HIV versus HIV. The Sirtuin Signaling Pathway presented the opposite effects, which is a predicted downregulation in TB-HIV versus HIV while all other comparisons predict upregulation. Interestingly, the T cell receptor signaling pathway was downregulated in mono-infections but upregulated during co-infection.

Due to the prominent disruptions predicted in the IL-17 signaling network, an expanded analysis of genes contributing to these outcomes was performed **(Fig 8).** Consistent with our previous results for decreased Th17 signaling during TB-HIV versus TB, when comparing exclusively these 2 groups by RNAseq, IPA demonstrated IL-17 signaling as the most down-regulated pathway, and in five of the 20 first affected pathways **(Fig 8A)**. Genes primarily affected in the pathway during TB were IL-17A, IL17RA, CXCL8, CXCL5, IL1B, CXCL3, and CCL2. Transcription of genes in the IL-17 pathway was either downregulated or not affected in the HIV versus non-infected differentially expressed genes. In contrast, a predominant upregulation was observed due to TB compared to non-infected in signaling downstream of the IL-17 receptor **(Sup. Fig 8)**. Interestingly, a comparison of TB-HIV versus HIV revealed an upregulation of IL-13, RGS16, CSF2, MMP9, HSP90AA1, SRSF1, and TNFSF15 that were not upregulated by TB or HIV alone. Analysis of Th differentiation between TB-HIV versus TB showed a mixed pattern of increased and decreased Th1 and Th2 differentiation signaling but predicted decreased Treg and Th17 differentiation (**Fig 8B**), consistent with **Fig 6**. IL-22 signaling decrease was observed in the TB-HIV versus TB comparison, consistent with our previous results (**Fig 5**), without IL-22 receptor involvement.

**Fig 8.**
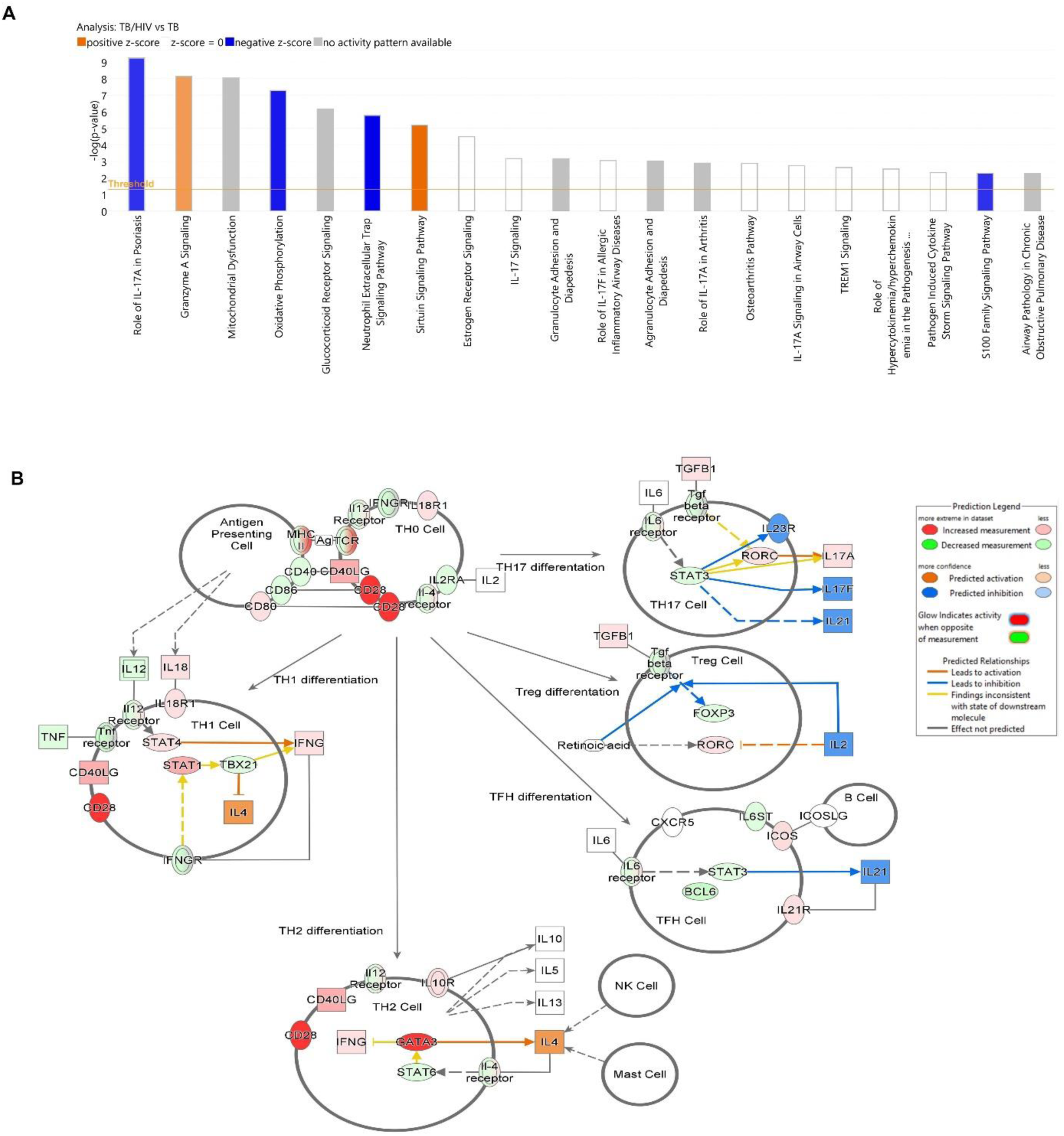
Transcriptional activation of Th17/IL-17 pathways due to Mtb infection of lung are impaired in Mtb-HIV co-infection. (A) Principal 20 canonical pathways differentially altered in comparison of TB-HIV versus TB, demonstrating enrichment of five pathways related to IL-17. Blue color depicts a negative activation z-score, predicting downregulation of the pathway. Orange color predicts upregulation by activation z-scores. (B) Th differentiation during TB-HIV versus TB comparison. Red and green show an increased or decreased experimental measurement respectively. Different orange shades show the confidence in predicted activation, while blue shades demonstrate confidence in predicted inhibition. Predictions were calculated through Ingenuity Pathway Analysis.

## 4. Discussion

Our findings in an HIS mouse model of acute TB and HIV co-infection provide an important advancement in our understanding of how HIV perturbs and exploits different Th subpopulations in the setting of pulmonary TB. To date, HIV targeting of Th subsets is poorly defined in tissue compartments beyond the gut. Similarly, the impact of HIV on depletion and functional immune responses in those with important co-morbidities such as TB are incompletely defined.

Our results (**Fig 1**) suggest that Th17 may be 1) an important and early target for HIV infection in the peripheral immune system, 2) a depletion-resistant niche for HIV replication at sites of Mtb-driven inflammation in the lung, and 3) an important mediator of effector function in lung that is compromised by HIV infection. Th22 cells, on the other hand, appear to be reduced due to HIV infection as associated with disruption of cytokine networks that may impair differentiation.

HIV was increased in the lung during early Mtb-HIV co-infection compared to the HIV group, while the spleen and liver presented similar viral loads to the HIV mono-infected group. Th cells depletion due to HIV has been observed in other organs such as gut-associated lymphoid tissue in earlier stages before depletion in blood (68), in accordance with our results. Increased viral loads presented only in lung during co-infection could be due to the timing in our experiment and increased effector T cell recruitment to the lung by 21 days post Mtb infection, as observed in standard mouse models (69). On the other hand, in bronchoalveolar lavage (BAL) of TB patients Th cells have been reported to express increased HIV co-receptor CCR5, which could lead to increased viral reservoirs and replication in the lung. Accordingly, CCR5 is downregulated by Mtb antigens in human splenic cells (70), which could impair HIV replication. Nevertheless, in more extended infection periods or different conditions, HIV replication during Mtb co-infection could be increased in other organs besides the lung (50).

Higher macrophage numbers and classical activation signaling pathways were observed during TB but not during TB-HIV. Antigen presentation by macrophages is essential for the control of mycobacterial growth (71). Macrophages also play vital roles during HIV infection establishment and dissemination (72) and are known as HIV viral reservoirs in the absence of T cells (73). Decreased macrophage percentages could explain increased pulmonary bacterial load during co-infection (50). The cause of decreased macrophages is elusive, although we could hypothesize it is not due to apoptosis, as HIV impairs macrophage apoptosis (74, 75). Of note, our immunofluorescent results identified CD4-negative IL-17 producers infected with HIV in Mtb-HIV co-infected granulomas. Macrophages can also produce IL-17 (76), suggesting that alveolar macrophages producing IL-17 could serve as potential viral reservoirs in the lung during co-infection. Another cause for increased bacterial burden in the lung could be the specific localization of HIV to the granuloma. It is important to note that, to the best of our knowledge, early entrance or localization of HIV in the lung has not been reported due to tissue restrictions. Here, we were able to image the early HIV localization in the lung, close to the granulomas in agreement with Foreman et al (67), but also in a peribronchial pattern during Mtb-HIV co-infection.

HIV is known to functionally impair Mtb-specific Th-responses, even before the development of detectable impairment of systemic immunity(77, 78). Th cells in the gut mucosa are depleted during primary HIV infection, 2-4 weeks post-exposure in human subjects (68, 79) and as soon as 7 days post-SIV infection in an NHP model (80). Reduction of lung interstitial Th cells has been reported at four weeks post-HIV infection in animal models (6), with loss of Th cells in TB granulomas as early as 14 days post-SIV infection in NHPs before depletion in blood is observed (67). Th17 are known to be preferentially depleted by HIV in human blood and different gastrointestinal regions (34–38, 40). However, preferential infection by HIV of Th cells in organs of Mtb-HIV co-infection, such as lung and spleen, is not defined due to understandable constraints in human sample acquisition. In a HIS model of post-drug TB relapse due to HIV, levels of pulmonary IL-17 were previously shown to be suppressed (50). Results of the current study demonstrate that Th17 are preferentially infected by HIV in the spleen, compared to other Th subsets. Interestingly, Th17 are infected with HIV at similar frequencies to Th1 and ThIL-10+ cells in the lung regardless of whether or not animals also have Mtb infection. This was a surprising outcome since we observed an increased viral burden in the lung of mice with co-infection.

Surprisingly, despite the increased Th17 percentages, their preferential infection in the spleen during HIV mono-infection, and their ability to harbor HIV in the lung granulomas, they were not preferentially infected in the lung or spleen during Mtb-HIV co-infection. Th17 from HIV-infected patients are not preferentially depleted in BAL in accordance with our results (36). However, BAL depletion of T cells does not reflect lung interstitium (6) and in the report of Foreman et al, Th1Th17 were preferentially infected in the granulomas and not in BAL, supporting a role for granuloma/localization-specific HIV responses (67). Another possible confounder for the lack of preferential infection of Th17 in the lung, is the early stage of co-infection, which may not yet reflect preferential infection or depletion of specific Th cell subsets. A decrease in polyfunctional Th1Th17 cells (or Th1*) is present in the Mtb/SIV co-infected granulomas (67). Even though Th17 were not preferentially infected in our results, future measurements of HIV-preferential infection could assess granulomas rather than total lung, polyfunctional Th1Th17 cells, and later stages of co-infection.

Despite lack of preferential pulmonary infection in our experiments, Th17/IL-17 signaling pathways were nonetheless affected due to Mtb-HIV co-infection. IL-17 signaling pathway genes (e.g., IL17RA, CXCL8 and CXCL5) and cytokine genes that regulate Th17/IL-17 bias (e.g. IL-6, IL-23, and IL-1β) were reduced during co-infection compared to TB alone. Due to the protective roles of IL-17 in TB (19, 20, 22–24), the loss of Th17-mediated IL-17 signaling is a previously underestimated risk for TB progression during Mtb-HIV co-infection, which potentially could not be rescued by other IL-17 producers, such as ILC3 (81), or CD8 T cells (5). However, as our differential transcriptome results reflects total lung, we cannot assess if the loss of IL-17 signaling originates from IL-17 producers other than Th17, such as macrophages, γδ T cells, or neutrophils (76).

Th22 are a considerably large T helper population in response to Mtb in lung and blood and are a major contributor to Mtb response during latent TB infection (LTBI) (29, 41, 44). However, Mtb-specific Th22 cells are markedly reduced in the blood during co-infection, even more than Th1 and Th17 (44) and are depleted by HIV in blood of LTBI individuals (41). Additionally, Th22 are depleted from blood and colorectal mucosa of SIV-mono-infected NHPs(40). We expand on Makatsa’s and Bunjun’s co-infection studies and show that Th22 are not only depleted from blood during Mtb-HIV co-infection (41, 44) but also in lungs and spleen. This reduction was significant compared to other T-cell subsets in HIV mono- and co-infection and was observed very early in the co-infection process. Makatsa (44) offers a postulate that HIV may deplete Th22 as these cells present enhanced HIV permissiveness and higher viral co-receptor (CXCR4 and CCR5) expression. Another potential mechanism observed in our results is the reduction of cytokines that promote Th22 (such as IL-6, IL-23, IL-1β and TNF). These cytokines were reduced during co-infection, which could impede differentiation of both Th22 and Th17 subsets.

Th22 are responsible for epithelial integrity and wound healing (17). Therefore, in the context of granuloma pathology for co-infection, the loss of Th22 could increase immunopathogenesis and TB progression. Interestingly, during mono-HIV/SIV infection other non-T-cell populations, such as ILCs, could rescue IL-22 production and maintain gut epithelial integrity, as demonstrated in the sigmoid colon (39, 81). Conversely, in Mtb mono-infection with CD4 depletion, IL-22 production could not be stably rescued by other cells (5). Furthermore, IL-22 rescue has not been demonstrated in the lung or during co-infection.

The lungs serve as a critical HIV viral reservoir, and despite ART, this organ presents increased persisting infectious and non-infectious diseases (82). Tissue reservoirs are an important reason why HIV infection remains incurable even with ART. Human lung parenchyma harbors an average of 6×10^9^ lymphocytes as potential HIV viral reservoirs (83, 84). Th17 have been described as a long-lived HIV-viral reservoir in the gut (57). Our results show that pulmonary Th17 cells were increased along with greater viral burden in the Mtb-HIV co-infected group, compared to HIV mono-infection, and frequently found to be infected with HIV in TB granulomas. Importantly, Th17 were the preferred Th target for HIV in the spleen. Together, these results could suggest Th17 as previously unforeseen viral reservoirs increased in number during co-infection.

The TB vaccination field currently aims to boost Th17 as a protective TB response as reviewed by (18, 64, 65, 85, 86), with some vaccines achieving it (24, 87–89). HIV infection can shift the Mtb-specific Th17IL-10+ “protective” responses to Th17Th1 “pathogenic” responses during Mtb-HIV co-infection (66). Our results demonstrate that Th17 are preferentially targeted by HIV in spleen and effectively infected in the lung. After boosting Th17 through vaccination, HIV could exploit an increased memory-Th17 population to promote viral pathogenesis, leading to increased viral reservoirs, suppressed cytokine responses, and ultimately vaccine failure. This poses interesting new questions about the Th17 role during co-infection that could lead us to a potential paradigm shift in vaccination efforts, primarily in high-burden Mtb-HIV co-infection countries(1).

In conclusion, we identified Th17 and Th22 impairments in the lung and spleen due to HIV that could further damage the antitubercular responses during co-infection. Th22 depletion due to HIV leads to decreased IL-22 during mono and co-infection. IL-22 replenishment during Mtb-HIV co-infection requires additional studies, as this cytokine is vital in wound healing and immunopathology prevention. Additional studies are needed to understand Th17 preferential infection by HIV in the lungs as it relates to experimental timing, polyfunctionality, and granuloma Th cell preference. Nevertheless, our studies did reveal decreases in IL-17 signaling pathways that could worsen TB pathogenesis. TB vaccination efforts should take into account these impaired responses for Mtb-HIV co-infected populations. Future vaccine efforts for HIV mono-infection should also consider the impairments in Th17 and Th22 cells in the lung and spleen.

## Acknowledgements

Y.B. Martinez-Martinez was supported by Conacyt-I2T2 in Mexico, Contex, and the James W. McLaughlin Fellowship Fund. Grant support was provided by NIH R01AI147948 award to J. Endsley. We thank the UTMB Flow Cytometry and Cell Sorting Core Facility, the Aerobiology Core Facility, and the Animal Resource Center for assistance in performing flow cytometry, aerosol infections with Mtb, and animal husbandry, respectively. We also thank the UTMB Optical Microscopy Core, the Next Generation Sequencing Core facility, and the UTMB Anatomical Pathology Laboratory for assisting in multifluorescent imaging, RNA sequencing, and histopathological services, respectively.

---

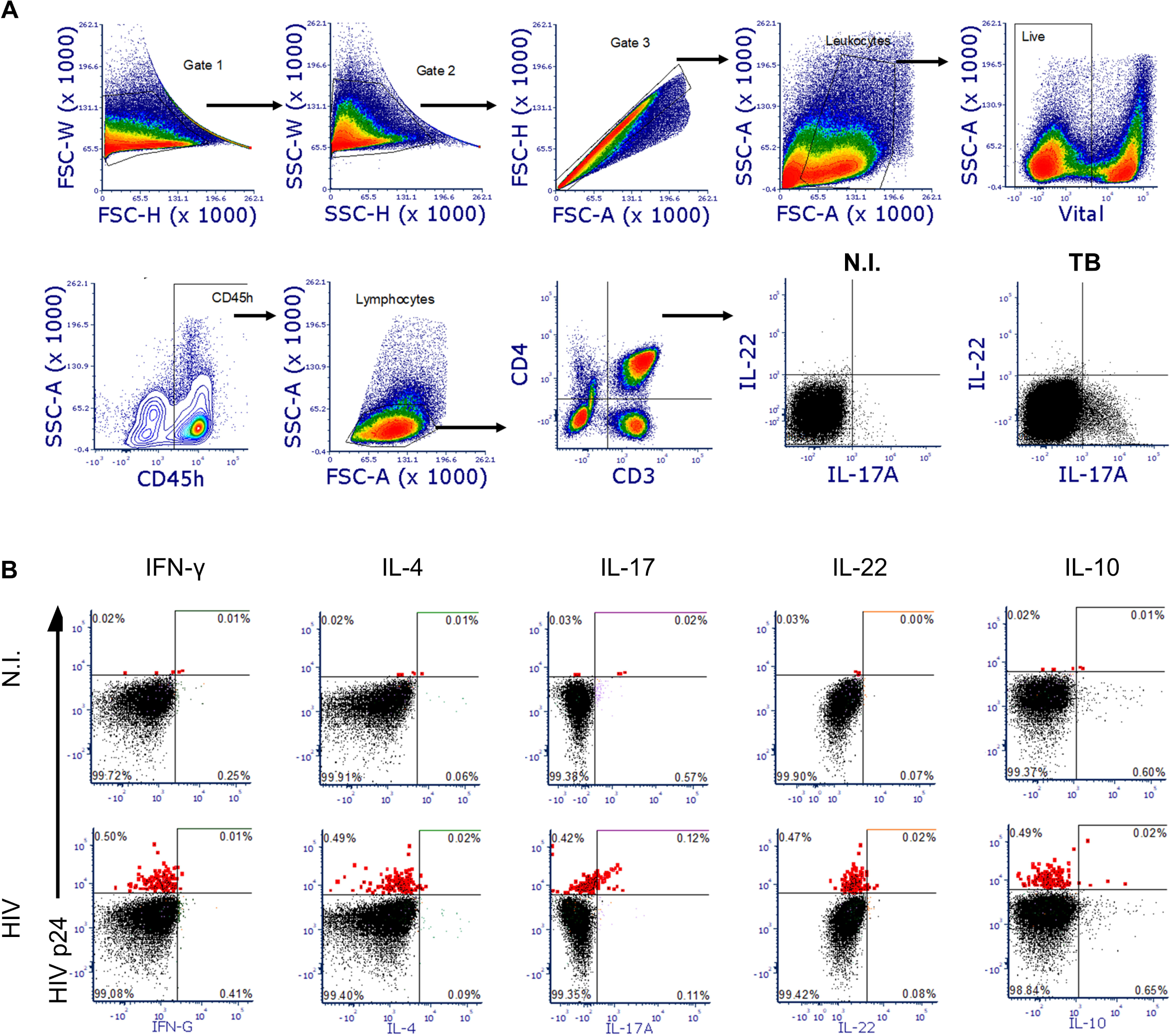

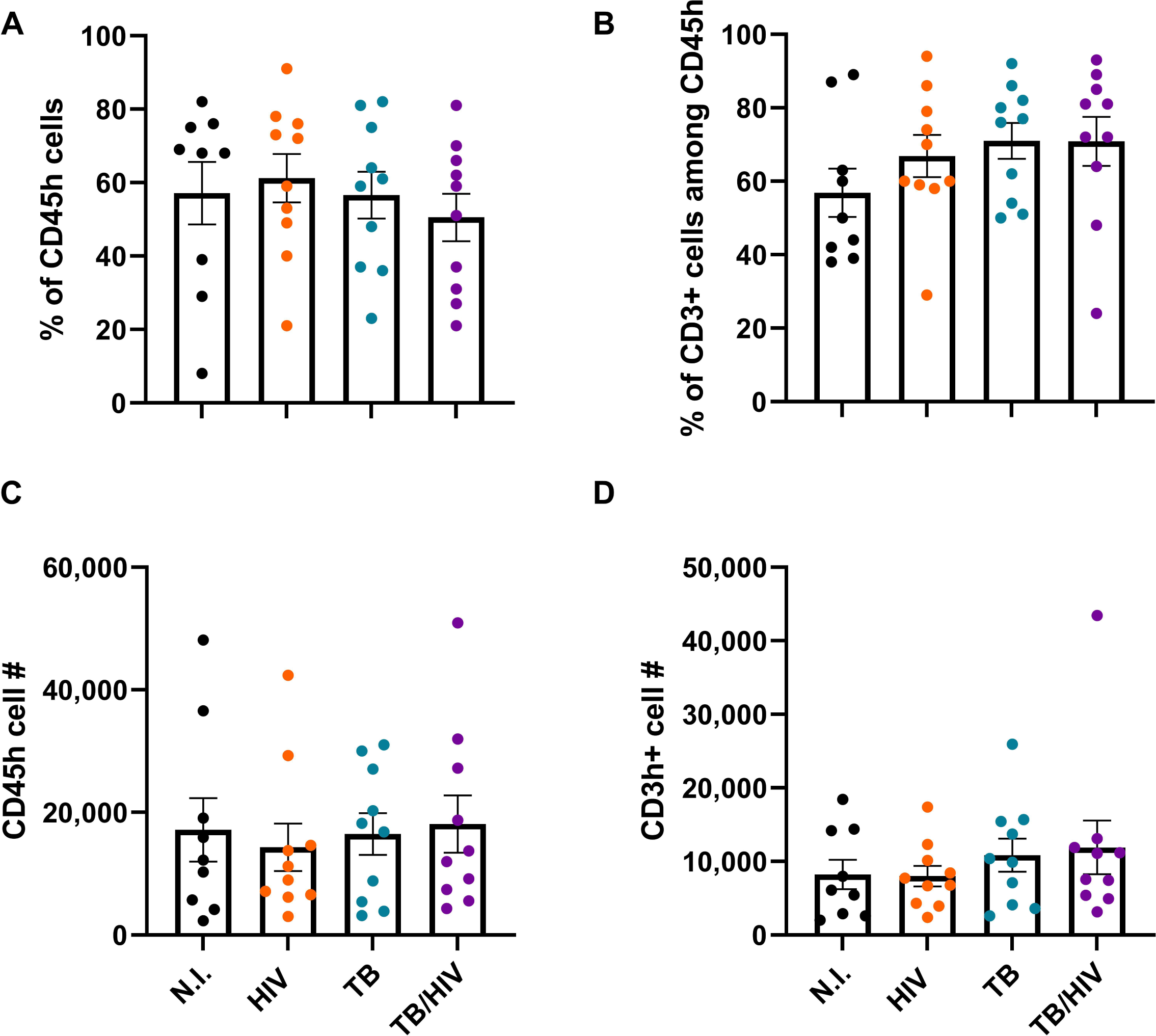

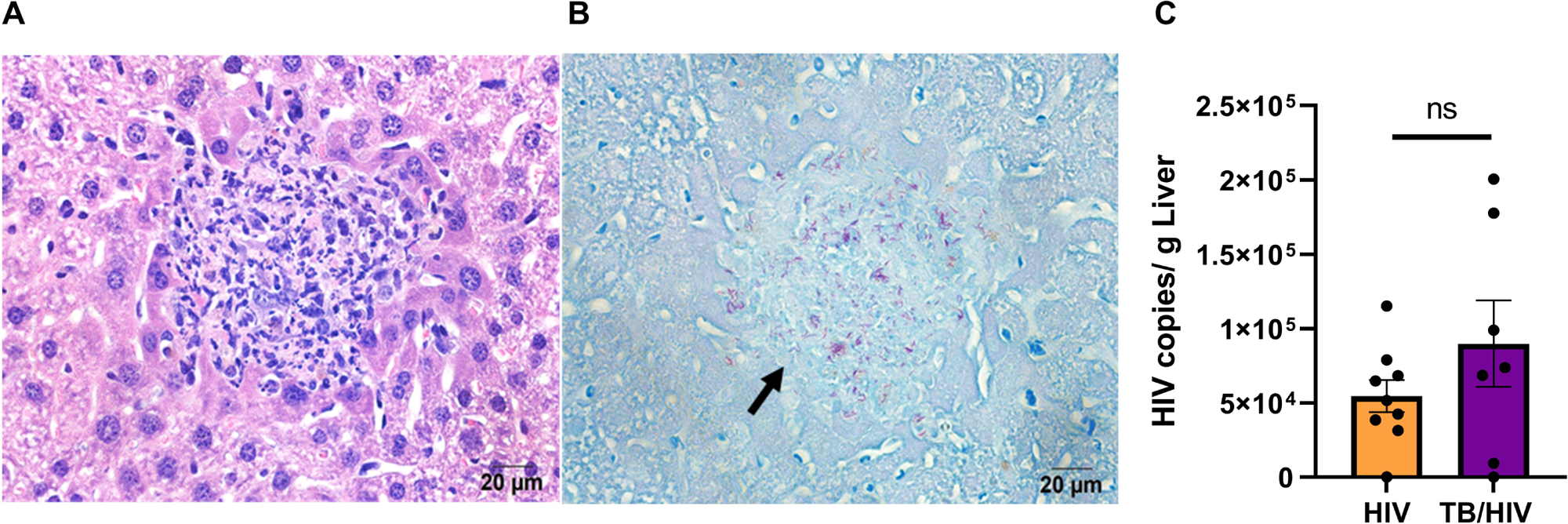

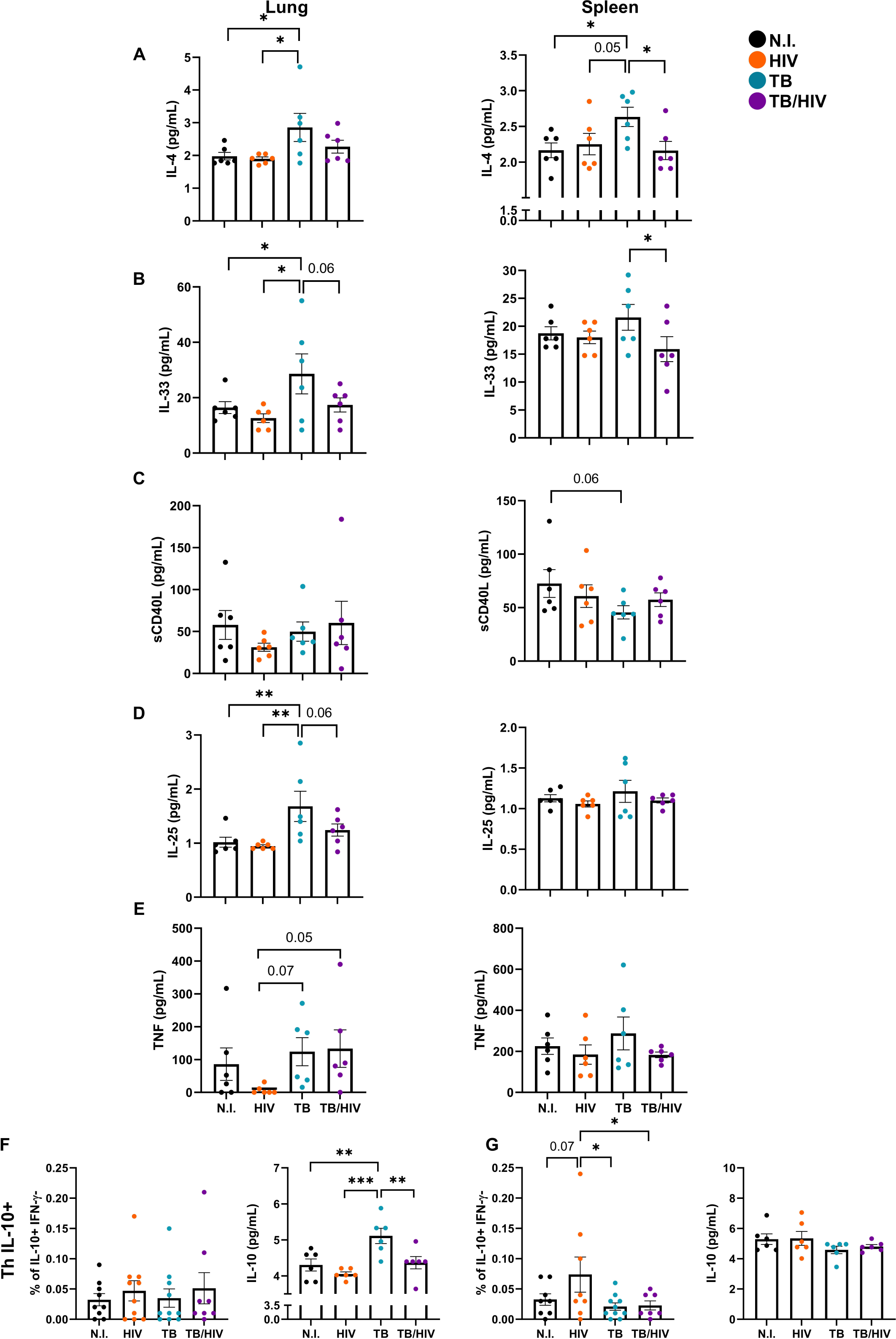

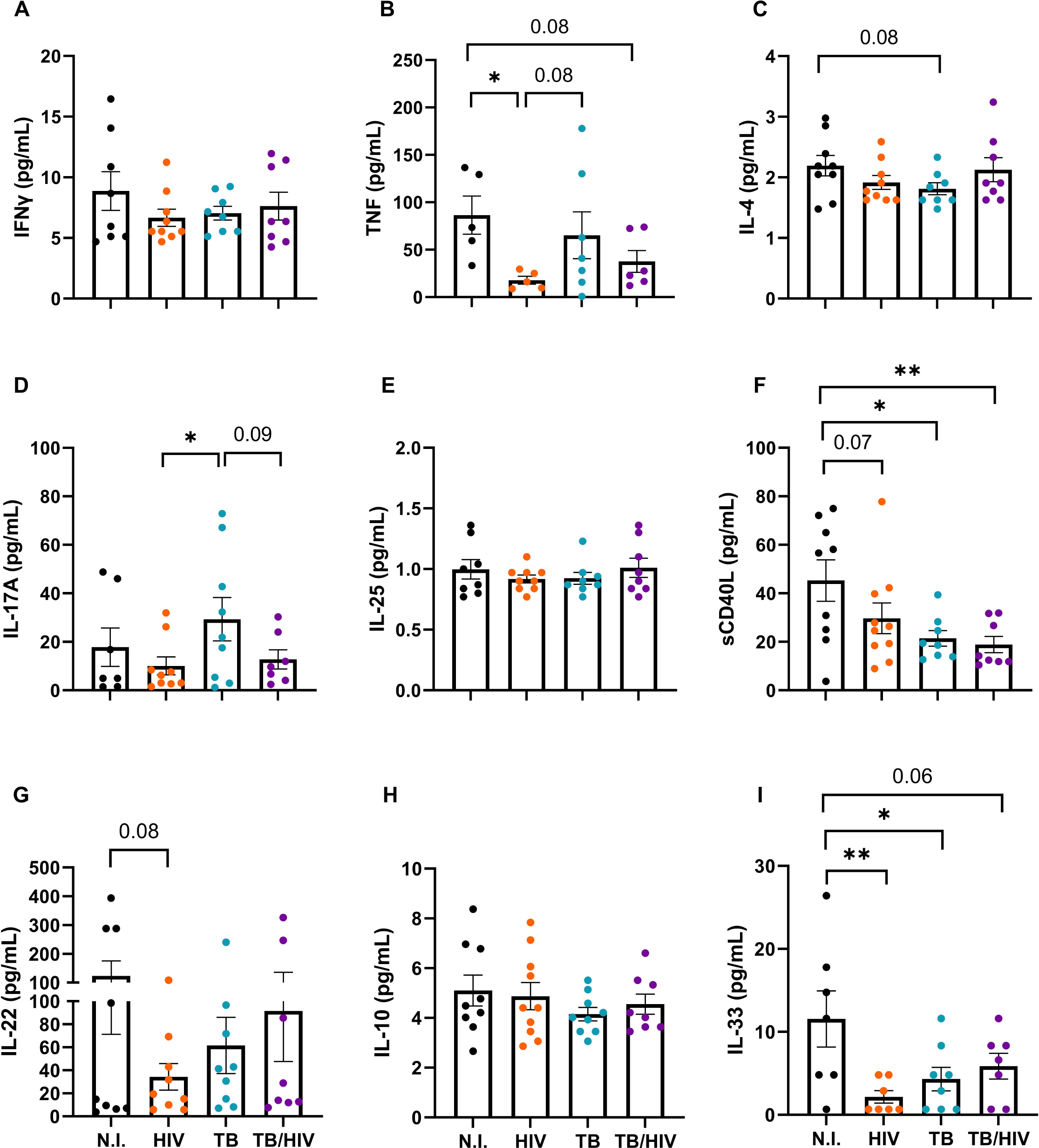

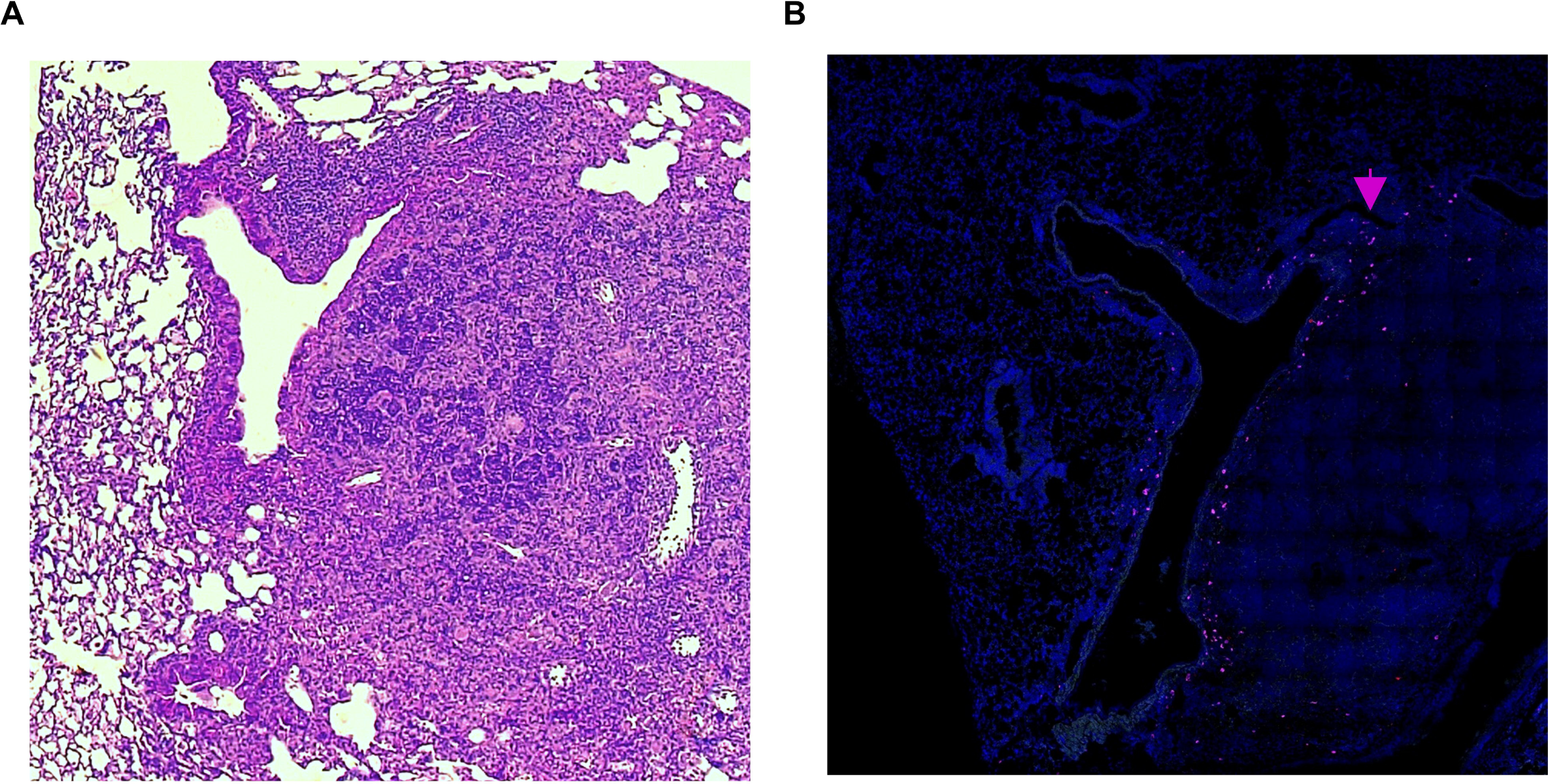

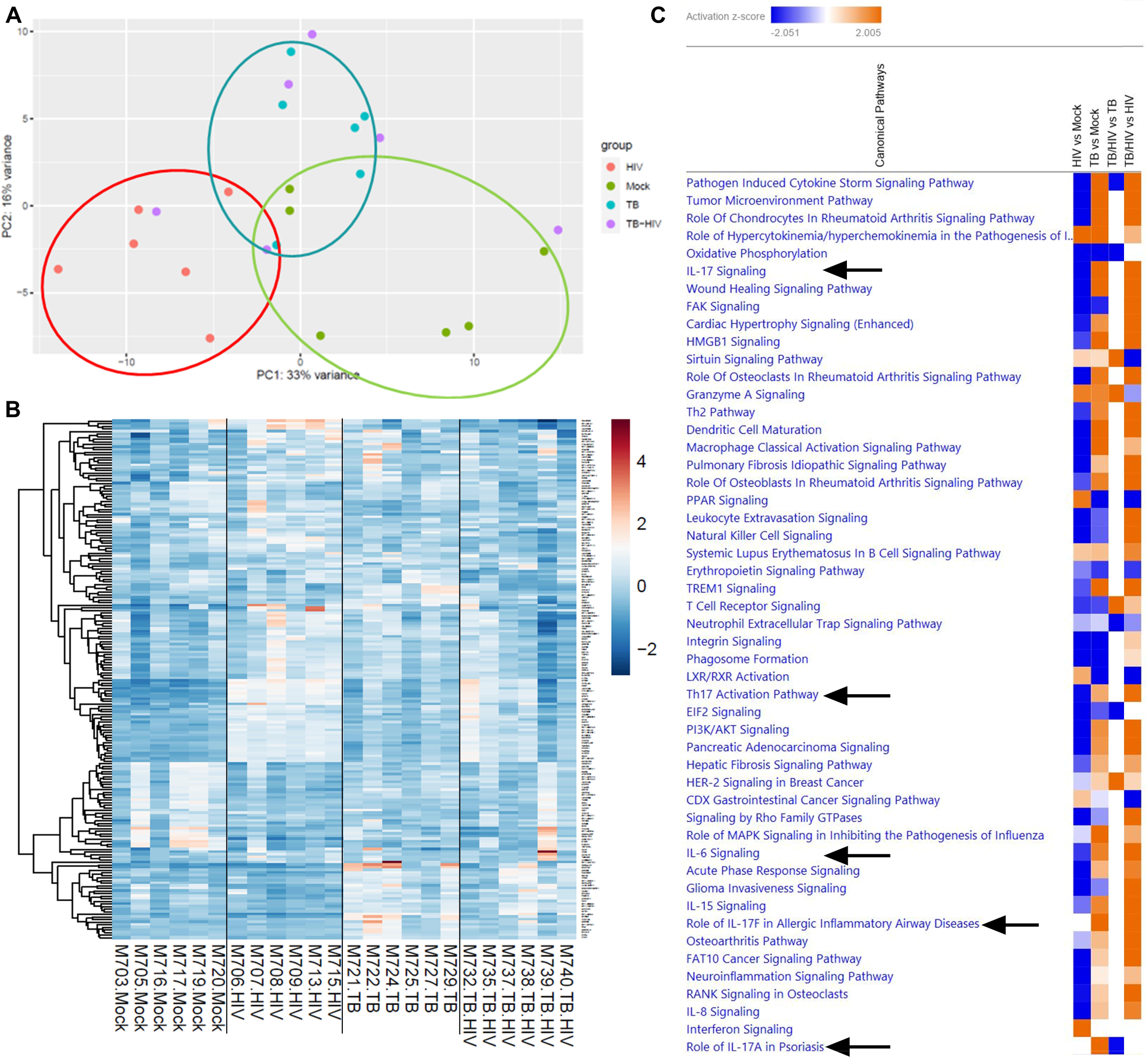

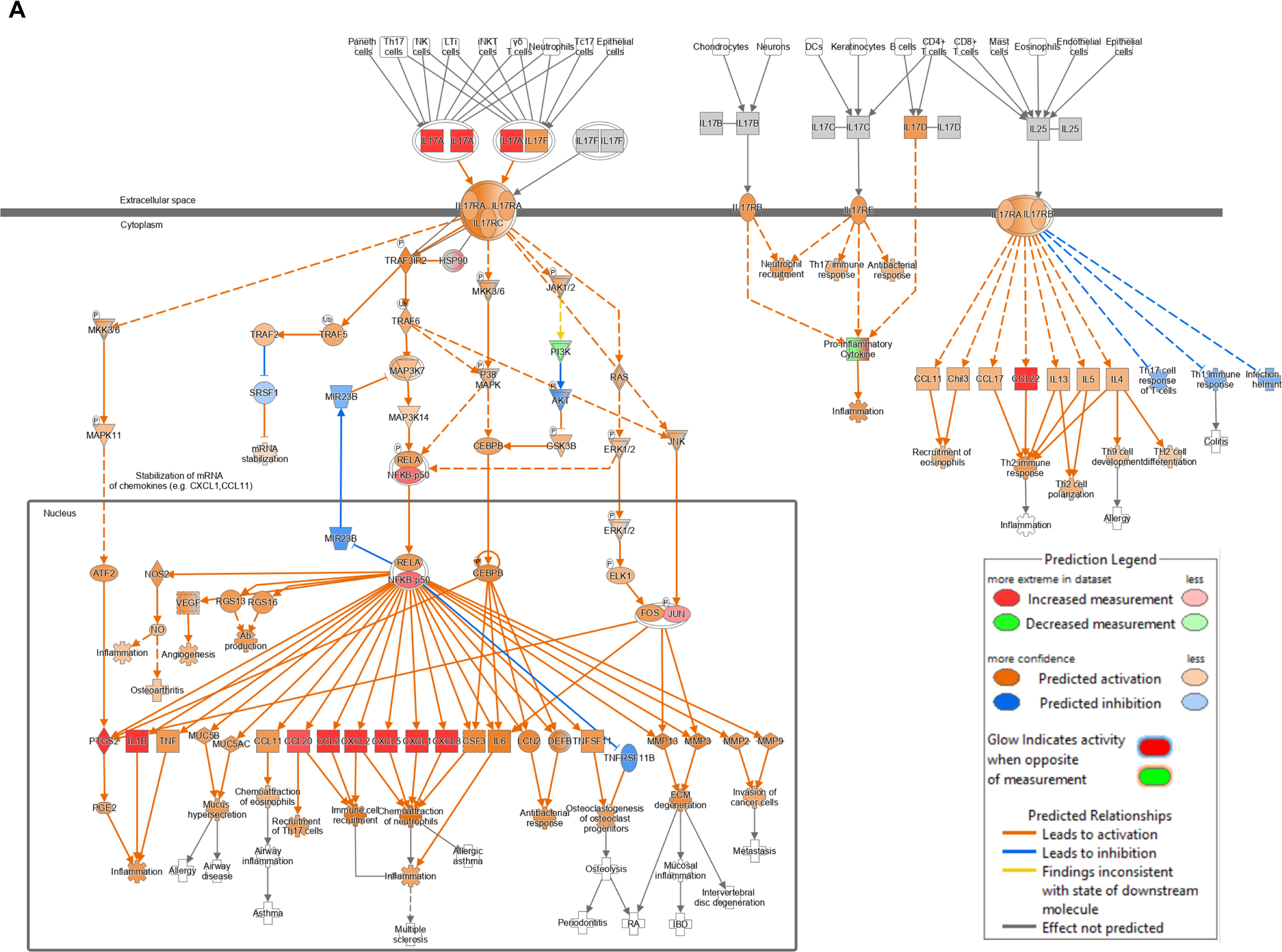

