## Supplementary figures for "HIV IMPAIRS AND EXPLOITS PULMONARY TH17 AND TH22 CELL-MEDIATED IMMUNE RESPONSES TO *MYCOBACTERIUM TUBERCULOSIS*"

**Running title:** Dysregulated Th17 and Th22 and Mtb-HIV co-infection

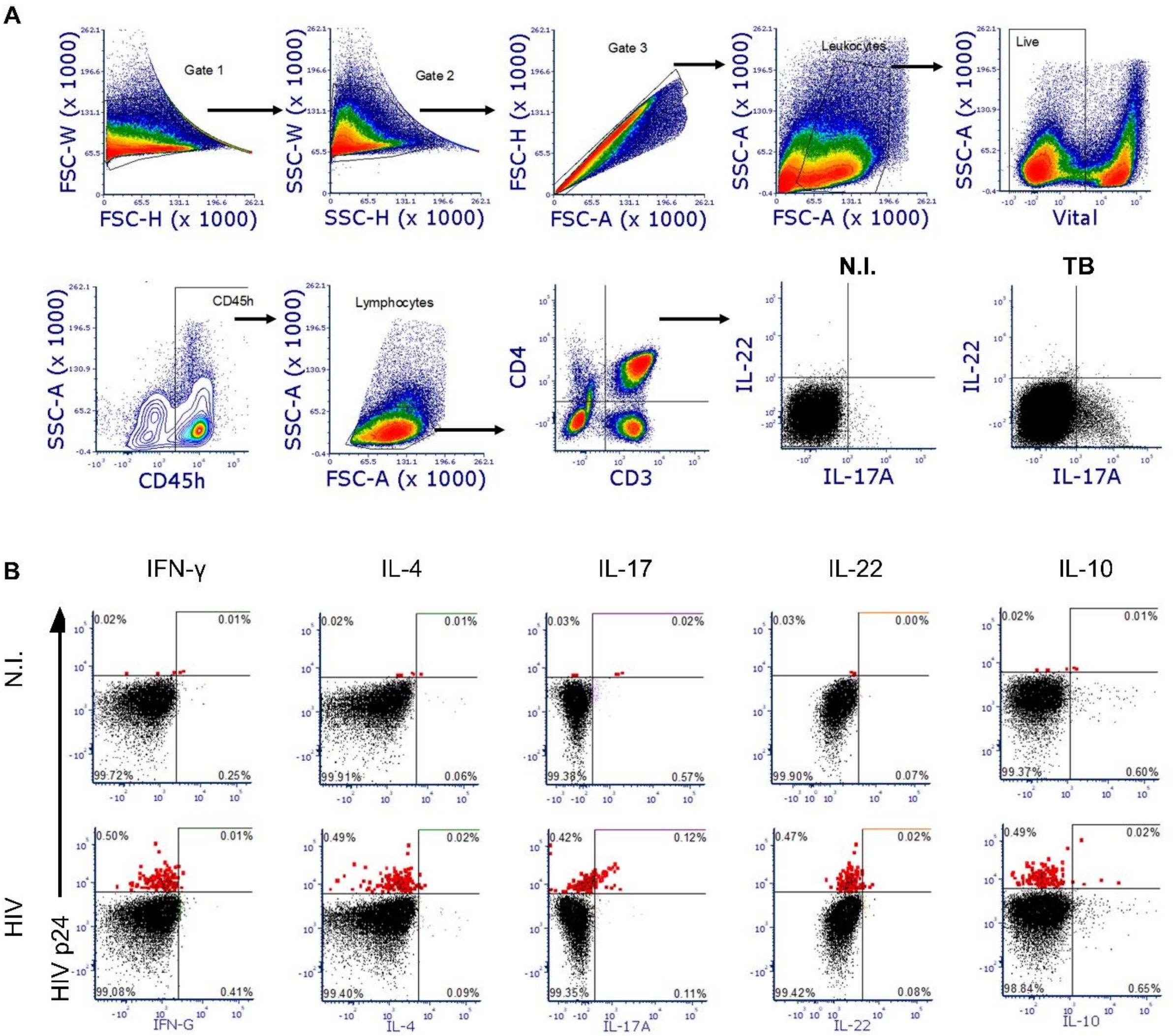


**Sup Fig 1. Representative flow cytometry gating strategy used for experiments.** (A) Gating used for selection of lung and spleen populations in both experiments. Sequential gating combinations of FSC-A, FSC-H, FSC-W, SSC-H, and SSC-W selected singlets. Among singlets (gate 3), Leukocytes were selected by size (FSC-A) and granularity (SSC-A). Viable (vital, negative for Live/dead Near IR) human CD45+ cells (leukocytes) were further gated for human macrophages (CD68+CD14-). Among leukocytes, lymphocytes were selected for T cells (CD3+), and T helper (CD3+CD4+) markers, which were additionally assessed for intracellular cytokines. In the example graphic, Th17 and Th22 are assessed by IL-17 and IL-22 intracellular markers, respectively, shown in the right panel of figure A within a representative uninfected and infected mouse. (B) Representative gating used in spleen for identification of HIV in the different cytokine producer subsets in figure 1E and H, among spleen controls (N.I., non-infected) (top) and HIV-infected spleen samples at 14 dpi (bottom).

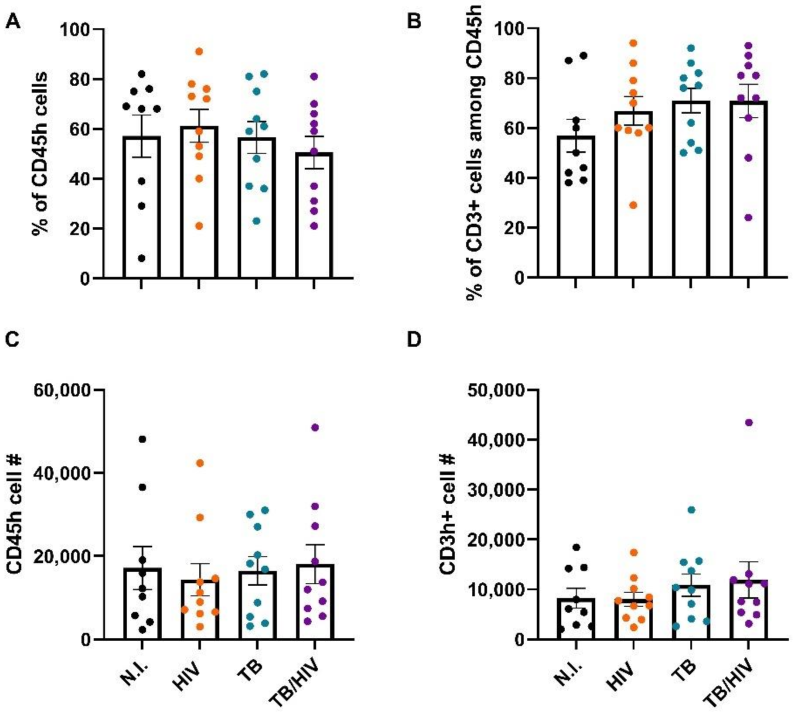


**Sup Fig 2. Reconstitution of human leukocytes in blood of HIS mice prior to initiation of co-infection experiment.** HIS mice were equally separated into four groups of 9-10 mice/group. Retroorbital blood was taken to assess distribution of human responses by flow cytometry before aerosol Mtb infection (or day 0) as in Figure 2A. (A) Reconstitution percentage, evaluated as % of human CD45+ (leukocyte) cells, among total leukocytes, using mouse and human CD45 markers. (B) Percentage of human T cells (CD3+), among the total human CD45+ leukocyte population. (C) Human CD45 leukocyte cell count. (D) Total human CD3+ cell count before infection among the four experimental groups: Non-infected, HIV, TB, and TB-HIV. No statistical difference was found before the infection (One-Way ANOVA with Benjamini FDR correction).

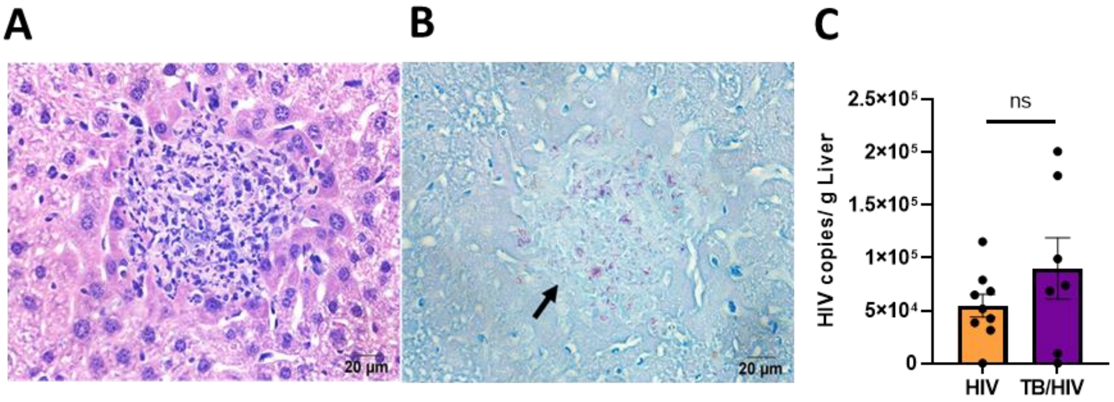


**Sup Fig 3. Liver histology and HIV infection in HIS mice.** (A) Representative Mtb infected H&E staining, showing a granulomatous lesion in liver. (B) Same histological area (arrow), with bacilli observed by AFB (magenta) staining. (C) HIV viral copies per gram of liver as detected by using RT-PCR. Ns= non-significant result based on Student's T-test).

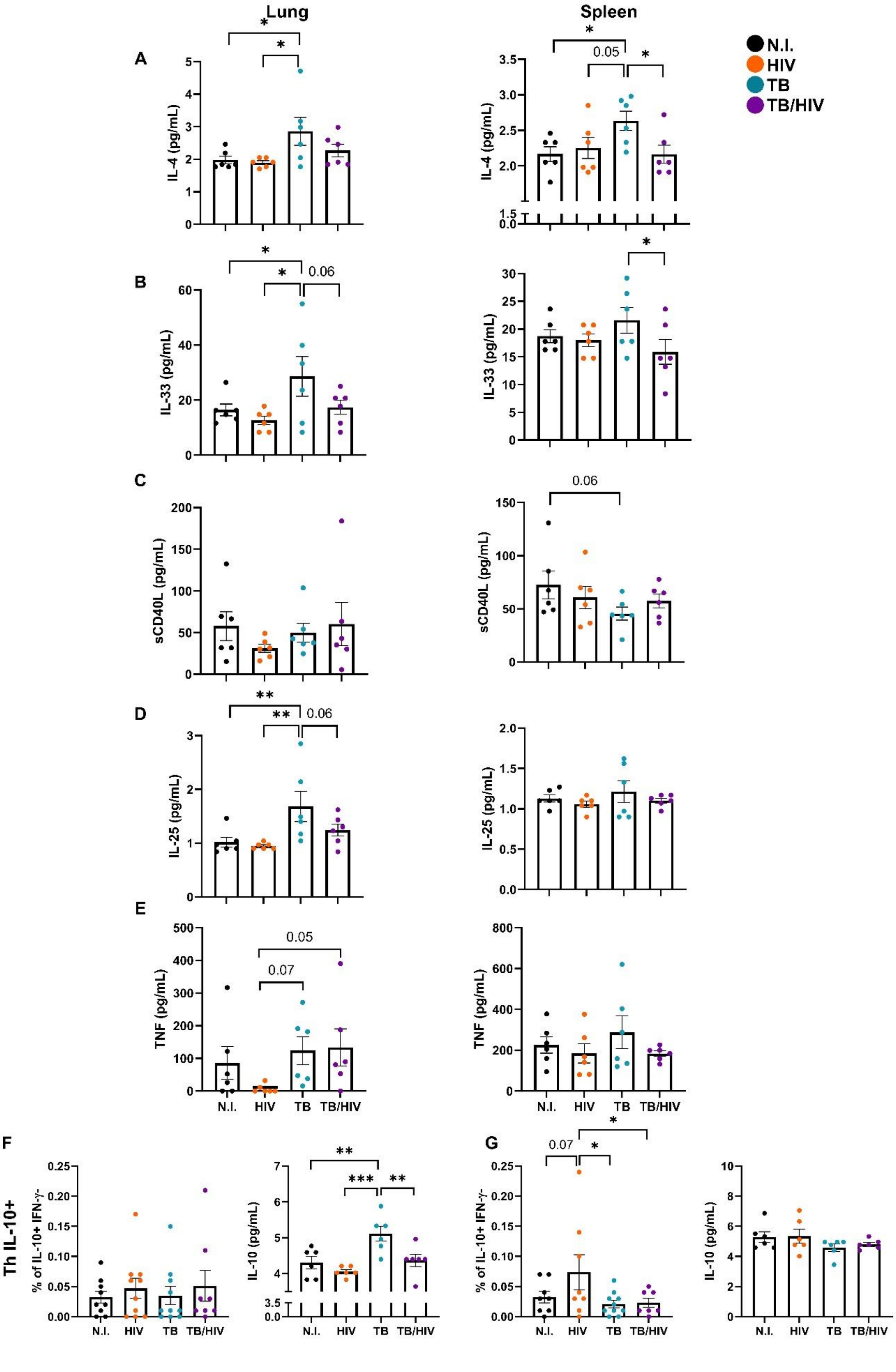

Lung
Spleen
N.I.
HIV
TB
TB/HIV
A
B
C
D
E
F
G
IL-4 (pg/mL)
IL-33 (pg/mL)
sCD40L (pg/mL)
IL-25 (pg/mL)
TNF (pg/mL)
Th IL-10+
% of IL-10+ IFN-γ-
IL-10 (pg/mL)
0.05
0.06
0.07

**Sup Fig 4. Soluble cytokines and ThIL-10+ cells in lung and spleen by infection status.** Supernatants were collected after cellular disaggregation for flow cytometry, frozen, and irradiated. After protein concentration, cytokines were quantified in supernatants as pg/mL by multiplex ELISA. In A-E we observe cytokines in lung (left panels) and spleen (right panels) including (A) IL-4, (B) IL-33, (C) soluble CD40 ligand, (D) IL-25 and (E) TNF in the four experimental groups: non-infected , HIV , TB , and TB-HIV groups. (F-G) Percentage of Th IL-10+ (IL-10+IFNγ-)(left), or IL-10 cytokine production (right) in lung (F) or spleen (G). *$p<0.05$, **$p<0.01$, ***$p<0.001$, (One-Way ANOVA with Benjamini FDR correction).

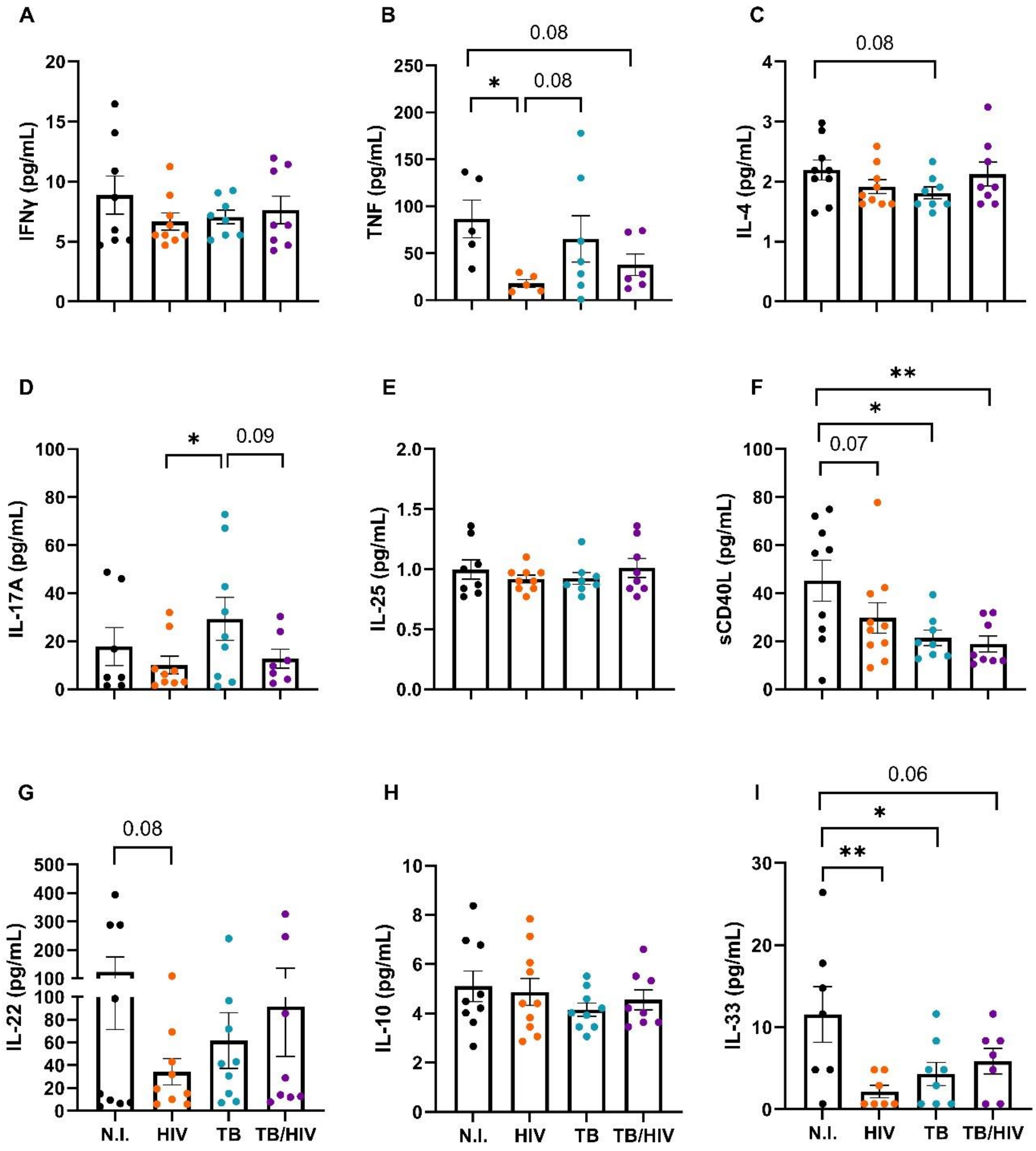


**Sup Fig 5. Human plasma cytokines produced during acute Mtb-HIV co-infection in the HIS mouse model.** Plasma samples were obtained after terminal intracardiac puncture, frozen, irradiated and cytokines were measured by multiplex ELISA. The different experimental groups are depicted with different color dots. From left to right, non-infected,, HIV , TB , and TB-HIV . Plasma cytokines quantified in pg/mL were IFNγ (A), TNF (B), IL-4 (C), IL-17A (D), IL-25 (E), soluble CD40 ligand (F), IL-22 (G), IL-10 (H) and IL-33 (I). *p<0.05, **p<0.01, (One-Way ANOVA with Benjamini FDR correction).

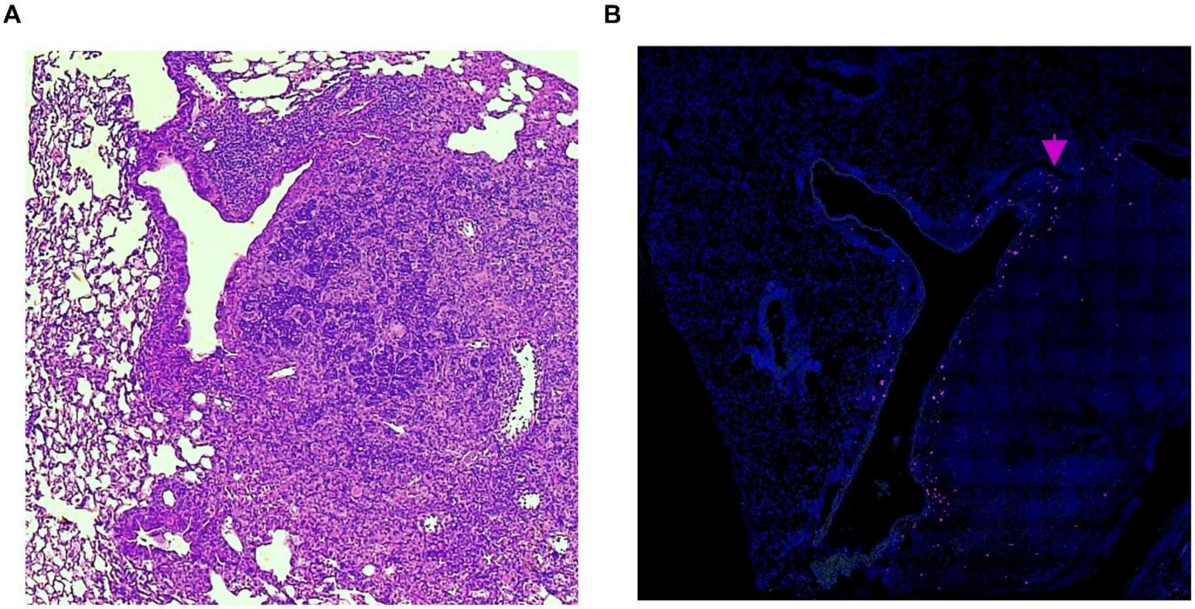


**Sup Fig 6. HIV+ cells co-localize to TB granulomas and peribronchial areas of the lung in the HIS mouse model of co-infection.** Organ scan overviews of lung in the TB-HIV group by H&E (A) or the respective tissue section RNAscope analysis (B), depicting HIV in cells surrounding the periphery of granulomas and the peribronchial areas of the lung by 9 days of HIV co-infection. In B, the representative immunofluorescent RNAscope image was stained through *in situ* hybridization, detecting RNA of HIV (non-gag-pol) with OPAL 690 (purple).

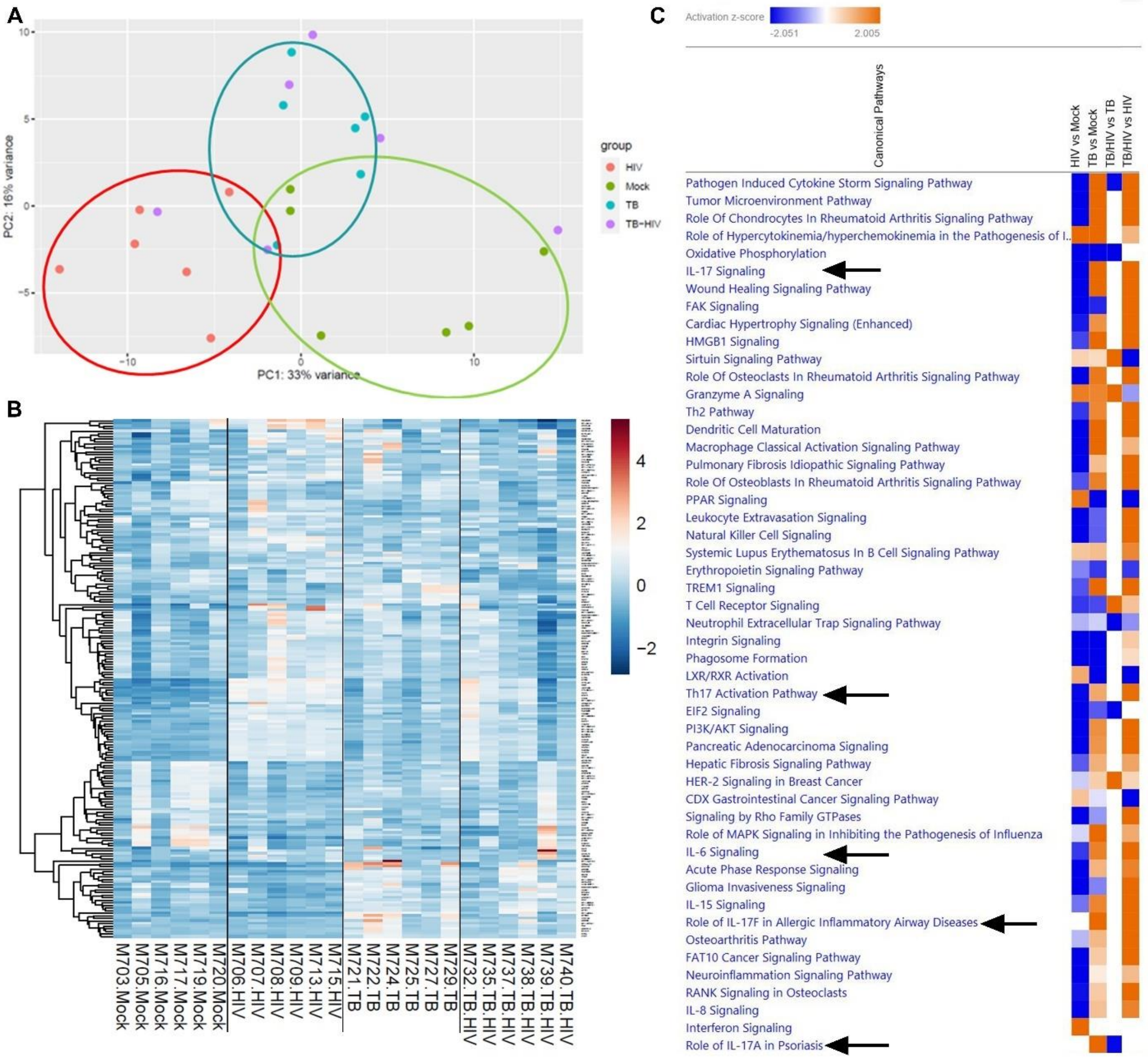


**Sup Fig 7. Human lung responses assessed by RNA sequencing demonstrates dysregulation of Th17/IL-17 pathways during Mtb-HIV co-infection in the HIS mouse model.** (A) Principal component analysis clusters the human lung responses according to infection in the HIS mouse model. Each dot represents an infected animal with HIV, Mtb, Mtb-HIV, or uninfected. (B) Heat map to observe the top 200 variable genes across all the samples, depicting transcriptome changes in the lungs. (C) Principal 50 canonical pathways affected by comparison of the four experimental groups, with arrows indicating IL-17 related pathways. The predictions were calculated through Ingenuity Pathway Analysis. Blue color depicts a negative activation z-score, predicting downregulation of the pathway. Orange color predicts upregulation by activation z-scores. n=6/group.

A

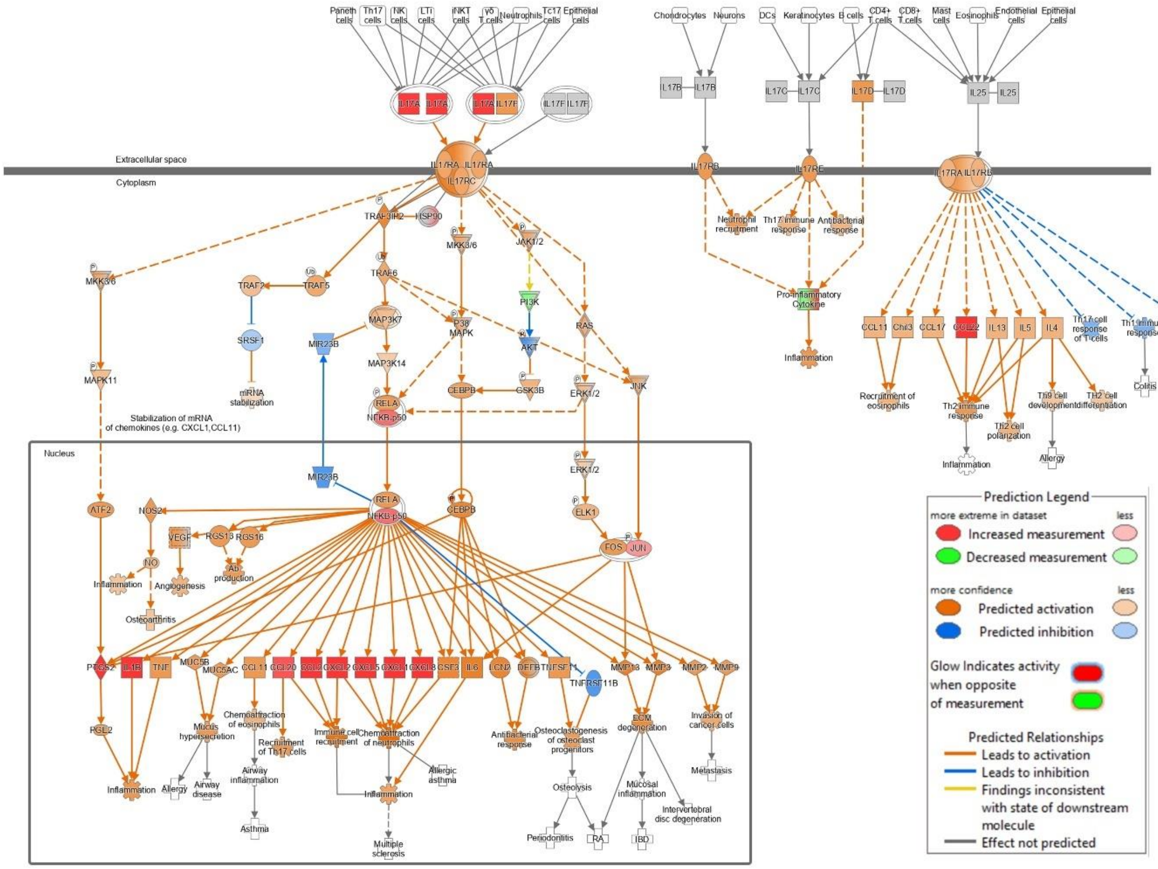


**Sup Fig 8. Differential transcriptome analysis identifies specific changes in the IL-17 signaling pathway.** (A) Signaling pathway comparison of TB versus non-infected. Most of the IL-17 signaling pathway is predicted to be upregulated (orange), due to experimentally upregulated genes (red), generated through Ingenuity Pathway Analysis.
